# Co-regulator activity of Mediator of DNA Damage Checkpoint 1 (MDC1) is associated with DNA repair dysfunction and PARP inhibitor sensitivity in lobular carcinoma of the breast

**DOI:** 10.1101/2023.10.29.564555

**Authors:** Joseph L. Sottnik, Madeleine T. Shackleford, Camryn S. Nesiba, Amanda L. Richer, Zoe Fleischmann, Jordan M. Swartz, Carmen E. Rowland, Maggie Musick, Rui Fu, Logan R. Myler, Patricia L. Opresko, Sanjana Mehrotra, Ethan S. Sokol, Jay R. Hesselberth, Jennifer R. Diamond, Matthew J. Sikora

## Abstract

Invasive lobular carcinoma of the breast (ILC) is typically estrogen receptor α (ER)-positive and presents with biomarkers of anti-estrogen sensitive disease, yet patients with ILC face particularly poor long-term outcomes with increased recurrence risk, suggesting endocrine response and ER function are unique in ILC. ER is co-regulated by the DNA repair protein Mediator of DNA Damage Checkpoint 1 (MDC1) specifically in ILC cells, driving distinct ER activity. However, this novel MDC1 activity is associated with dysfunctional canonical DNA repair activity by MDC1, but without typical features of DNA repair deficiency. To understand reciprocal activities of MDC1, we profiled the MDC1 interactome and found MDC1-associated proteins in ILC cells mirror a “BRCA-like” state lacking key homologous recombination (HR) proteins, consistent with HR dysfunction but distinct from classic “BRCAness”. HR dysfunction in ILC cells is supported by single-cell transcriptome and DNA repair activity analyses, with DNA repair signaling and functional data, showing dysfunctional induction and resolution of HR. In parallel, ILC tumor data are consistent with a distinct form of HR dysfunction via impaired HR resolution, lacking BRCA-like genomic scarring but showing elevated signatures of PARP inhibitor sensitivity. We demonstrate this HR dysfunction can be exploited using PARP inhibition, and found that talazoparib treatment produced a durable growth suppression both *in vitro* and in multiple ILC xenografts *in vivo*. ILC-specific ER:MDC1 activity creates a new context for ER and MDC1 function in ILC, at the cost of a DNA repair dysfunction, which may be therapeutically targetable.

**Significance:** ILC is rarely associated with biomarkers of HR deficiency, and as such patients are rarely eligible for treatment with PARP inhibitors. Our work suggests ILC presents with a previously unappreciated form of HR dysfunction, linked to ILC-specific genomic activity of ER and MDC1, which imparts sensitivity to PARP inhibition.

## INTRODUCTION

Invasive lobular carcinoma of the breast (ILC) is increasingly recognized as a distinct clinical and molecular entity among breast cancers (1–3). Beyond ILC’s unique histopathological presentation – i.e., infiltrating the breast in a discohesive, “single-file” pattern – we now appreciate that ILC have distinct genetic and cellular features relative to breast cancers of no special type (also known as invasive ductal carcinoma, IDC). Most ILC present with biomarkers of low-risk disease, as ∼95% are estrogen receptor α (ER)-positive and ∼80% are classified as the Luminal A molecular subtype (4), but retrospective studies suggest that ILC carry more long-term risk of poor outcomes than IDC. Retrospective studies show that 5-8 years after diagnosis, ILC recurrences increase relative to IDC regardless of ER status, with ER+ ILC having a ∼30% increased risk of recurrence vs ER+ IDC (5–8). Further, recurrent or metastatic ILC presents distinct challenges, as ILC are unique among breast cancers in metastasizing to the GI/GU, peritoneum, ovary/uterus, the orbit, and the leptomeninges (2,9,10). Treating ILC is further complicated due to controversies related to anti-estrogen resistance (11), the limited benefit of chemotherapy for most patients with ILC (2,12,13), and the lack of any approved precision or targeted treatments designed for ILC. New treatments for patients with ILC are needed, but strategies must be driven by advances in defining the unique biology of ILC.

We previously profiled the ER interactome in ILC cells with the goal of defining novel ER co-regulators that mediate ILC-specific endocrine response or anti-estrogen resistance (14). This approach identified Mediator of DNA Damage Checkpoint 1 (MDC1) as a critical ER co-regulator specifically in ILC cells. We found that the ER:MDC1 interaction is likely unique to ILC cells, MDC1 was specifically required for ER- driven growth in ILC but not IDC cells, and MDC1 knockdown broadly dysregulated the ER transcriptome in ILC cells. As MDC1 has no known enzymatic activity, the mechanism by which MDC1 co-regulates ER is unknown. The canonical role of MDC1, however, is to bind newly formed phosphorylated histone H2AX (γH2AX) marks at DNA double-strand breaks (DSB) and scaffold double-strand break repair machinery, including the MRE11/RAD50/NBS1 (MRN) complex and additional downstream partners that ultimately facilitate repair (15–17). The MDC1 scaffold initiates signaling via CHK1/2 to cause cell cycle arrest, and coordinates DSB repair either via homologous recombination (HR) or non-homologous end joining (NHEJ).

While we confirmed these canonical DNA repair roles of MDC1 in IDC cells, this was not the case in ILC cells. For example, γH2AX foci formation, which requires MDC1 (18), was limited in ILC cells and γH2AX instead presented largely as an elevated pan-nuclear signal, a pattern linked to DNA repair dysfunction (19). These observations collectively suggest that the novel ER co-regulator activity of MDC1 may come at a cost of compromised MDC1 activity in DNA repair.

ILC do not typically present with biomarkers of overt DNA repair deficiency such as germline *BRCA1/2* mutations. These mutations are uncommon in ILC and are depleted relative to IDC. ILC accounts for up to 15% of all breast cancers, but only ∼2% and ∼8% of breast cancers in *BRCA1* or *BRCA2* mutation carriers, respectively, and germline *PALB2* and *TP53* mutations are also rare in ILC (20–22). Similarly, somatic mutations in genes associated with DNA repair are uncommon in primary ILC, e.g. somatic *TP53* mutations were found in only ∼8% of ILC, and only 5.7% of Luminal A ILC, in the Cancer Genome Atlas (TCGA)(4).

ILC genomes are typically considered ‘quiet’ and overall show fewer structural re-arrangements than other breast cancers (8). Similarly, HR deficiency as defined by various mutational signatures is not common in ILC but is typically associated with the few ILC that are *TP53*-mutant or with germline *BRCA1/2* mutations (3).

Together these observations would suggest that DNA repair deficiency is uncommon in ILC. However, data from TCGA show that a putative transcriptional subtype of high-risk ILC, i.e. “Proliferative” ILC, was associated with an elevated DNA damage response (DDR) signature in protein array analyses (4). This is paralleled by frequent *MDM4* amplification in ILC tumors (17%) (23), suggesting suppression of p53-driven pathways by mechanisms independent of *TP53* mutation may be an underappreciated feature of ILC. Additionally, in Foundation Medicine data (FoundationCORE), primary ILC were shown to have increased tumor mutational burden relative to IDC, which further progress with metastatic ILC (24). Among metastatic ER+ breast cancers, 8.9% of ILC were TMB-high or extreme versus 1.5% of IDC (p=0.0016); ∼37% of ILC were classified as TMB-medium or higher versus only 25% of IDC (p=0.013) (24). Collectively these data suggest that our understanding of DNA repair signaling and function in ILC is incomplete, and may not be adequately defined as ‘proficient’ versus ‘deficient’ as it relates to *BRCA1/2* mutant tumors.

Based on the novel ER:MDC1 interaction in ILC cells and putative link to DNA repair dysfunction, we hypothesized that profiling the MDC1 interactome in ILC cells would both shed light on mechanisms by which MDC1 acts as an ER co-regulator, and offer insight into DNA repair function in ILC cells. These studies confirmed connections between the co-regulator activity versus DNA repair functions of MDC1, and identified potential mechanistic underpinnings of a “BRCA-like” phenotype in ILC cells. We explored whether MDC1- related “molecular BRCAness” in ILC cells provided opportunities for synthetic lethal therapeutic approaches. Better understanding of the cross-talk linking ER:MDC1, transcriptional regulation and DNA repair capacity offers unique opportunities to understand ILC biology and develop precision treatments around synthetic vulnerabilities unique to ILC.

## RESULTS

### The MDC1 interactome in ILC cells indicates DNA damage response dysfunction

We profiled the MDC1 interactome in ILC cells to identify transcriptional co-regulator partners of ER:MDC1, and to assess differential MDC1 association with canonical DNA repair partners, by performing MDC1 co-immunoprecipitation + mass spectrometry (IP/MS) in ILC versus IDC cells. Of note, proteomic approaches have shown MDC1 association with the MRN complex and other DNA repair proteins, but studies to date have used exogenous and/or tagged versions of MDC1 or MDC1 fragments, and only in HeLa, HEK293 (293), or U2OS cells (25–29). We performed endogenous MDC1 IP from ER+ cell lines MCF7 (IDC), HCC1428 (IDC, *BRCA2*-mutant), MM134 (ILC), and 44PE (ILC), as well as 293FT cells (to compare our approach to literature data). IP/MS was performed absent DNA damaging agents, from biological duplicate MDC1 IP samples and a matched IgG control per cell line. MDC1 was strongly enriched in targeted IP versus IgG across all samples (MDC1 iBAQ: ∼90-fold up to >1000-fold for MDC1 IP vs IgG, **Supplemental Figure 1A**).

Across all 5 cell lines, over 2000 proteins were enriched relative to IgG controls (**Supplemental File 1**). Identified proteins were most enriched for ribosomal and related mRNA/rRNA-processing proteins (as by Salifou et al (27), discussed below). We excluded/filtered RNA processing proteins to facilitate further analysis (see Methods), yielding n=1666 proteins identified in at least one cell line (**Supplemental File 1**). We first examined MDC1-associated proteins in 293FT cells (n=1497, n=1236 after filtering), and confirmed that the MRN complex (MRE11, RAD50, NBN), constitutive partners of MDC1 in 293FT and other cells, were strongly enriched (**Supplemental Figure 1B**). More broadly, DNA repair pathway proteins and epigenomic proteins were strongly enriched, the latter consistent with functions of epigenomic proteins in DNA repair and reported roles for MDC1 in chromatin remodeling (30) (**Supplemental Figure 1C, Supplemental File 1**).

We compared our 293FT data to MDC1 interactome data from Salifou et al (27), the only other proteome-wide MDC1 interactome data published to date. Whereas we utilized an MDC1-specific antibody for IP, Salifou et al used CRISPR/Cas9 to append a Flag-hemagglutinin (HA) tag to the N-terminus of endogenous MDC1 in 293FT cells, then performed tandem affinity purification of Flag/HA-MDC1, reporting n=384 MDC1 associated proteins (n=276 after filtering as above). As noted above, both datasets were strongly enriched for splicing and translational proteins (**Supplemental Figure 2A**), supporting a role for MDC1 in protecting genomic integrity at actively transcribed sites (27,31). Upon filtering, our data using direct MDC1 IP was specifically enriched in DNA repair and chromatin remodeling proteins (**Supplemental Figure 2B**). Of note, our data included greater depth of coverage of similar biological pathways compared to the Flag/HA-MDC1 dataset (**Supplemental Figure 2C**), and proteins in pathways enriched only in the Flag/HA-MDC1 dataset were still identified in our data (**Supplemental Figure 2D**). Direct MDC1 IP with MS confirms MDC1 association with mRNA/translational machinery and provides greater representation and enrichment for MDC1-associated DNA repair and chromatin remodeling proteins.

In the breast cancer cell lines, >700 MDC1-associated proteins were identified per cell line after filtering (**Figure 1A**). Hierarchical clustering showed that the MDC1 interactome in ILC cells was distinct from MCF7 (IDC) cells and more closely related to the *BRCA2*-mutant HCC1428 cells (**Figure 1A**), which was consistent with a relative depletion of MDC1-associated DNA repair proteins. DNA repair proteins were enriched overall in all 4 lines (**Supplemental File 1**); however, HR proteins were enriched only in MCF7 but depleted in the ILC and *BRCA2*-mutant cells (**Figure 1B**). This depletion of HR proteins included minimal/absent MRN complex proteins and other HR factors (**Figure 1C**), yet the MDC1 interactome in ILC cells was strongly enriched for epigenomic partners (**Figure 1D**), e.g. histone acetylation proteins. Distinct subsets of histone acetylation factors were identified in MCF7/IDC vs ILC (and *BRCA2*-mutant) cells, e.g. RUVBL1/2 in the former and the HBO1-complex (ING5, BRPF3, MEAF6, JADE3) in the latter (**Figure 1E**). Collectively these data suggest that the constitutive association of MRN with MDC1 may not be absolute, and that the absence/depletion of MRN on MDC1 is associated with DNA repair/HR dysfunction. The depletion of MRN and other HR proteins among MDC1-associated proteins in ILC is consistent with the pan-nuclear γH2AX localization we previously reported (14). However, the epigenomic functions of MDC1 likely remain intact despite potential DDR dysfunction, i.e. the pan-nuclear γH2AX signal (14) and lack of HR protein associations, and thus may be associated with the co-regulator functions of MDC1 in ILC cells.

**Figure 1.**
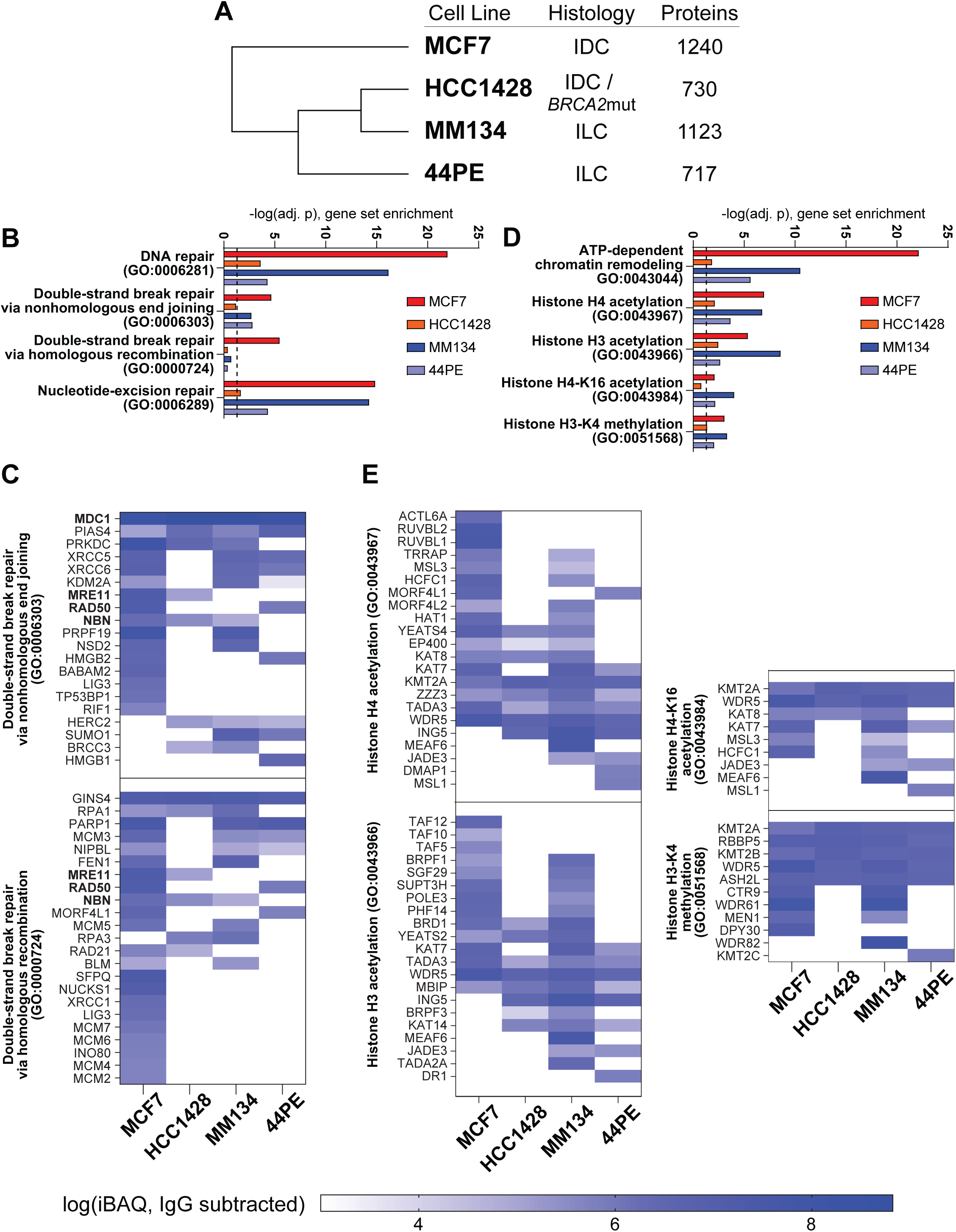
MDC1 interactome in ILC cells mirrors a BRCA-mutant state. **(A)** Hierarchical clustering (Pearson) of MDC1 associated proteins after filtering of RNA-processing related proteins. **(B,D)** Gene set enrichment against GO Biological Processes. Dashed line: adj. p = 0.05. **(C,E)** Mean IgG-subtracted intensity of duplicate IP/MS samples for proteins in indicated ontology groups.

### Inhibiting estrogen receptor in ILC cells restores DDR/HR proteins in the MDC1 interactome

We hypothesized that the novel ER:MDC1 interaction in ILC cells is at least in part responsible for remodeling the MDC1 interactome, and tested this by treating MM134 cells with the ER antagonist and degrader fulvestrant for 24 hours prior to MDC1 IP/MS. Hierarchical clustering showed that with fulvestrant the MDC1 interactome in MM134 more closely resembled MCF7 cells (i.e. HR-competent cells; **Figure 2A**). Strikingly, this change was associated with increased MDC1-association with HR pathway proteins (**Figure 2B**) including the MRN complex, specifically MRE11 and RAD50 (**Figure 2C**). Notably, HR proteins gained in MM134+Fulv versus MM134 (**Figure 2C**, orange bar) were all identified as MDC1-associated in MCF7 cells, suggesting that these MDC1 interactions were restored with fulvestrant treatment, rather than fulvestrant driving new MDC1 interactions. MDC1 association with epigenomic proteins remained enriched in fulvestrant treated cells (**Figure 2D**), but among histone acetylation proteins, several were shifted by fulvestrant treatment to resemble the state in MCF7 cells, including restoration of RUVBL2 and depletion of HBO1-complex proteins ING5, JADE3, and BRPF3 (**Figure 2E**, orange bar). These observations support that ER plays a role in regulating the cross-talk between genomic functions of MDC1 and the DDR functions of MDC1 at least in part through control over the MDC1 interactome.

**Figure 2.**
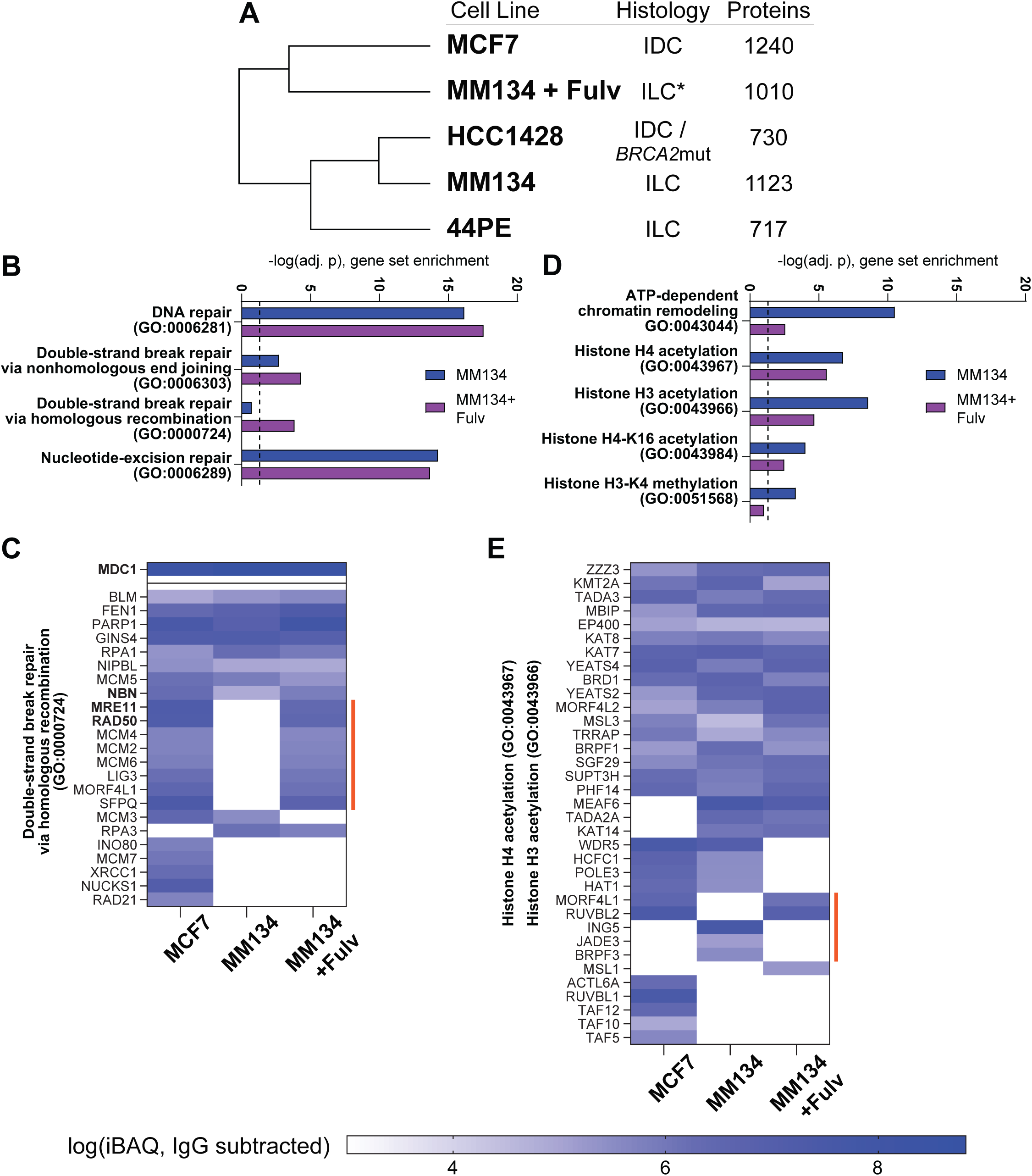
ER inhibition with fulvestrant shifts MDC1 interactome toward a non-BCRA-mutant state. **(A)** Hierarchical clustering (Pearson) of MDC1 associated proteins as in Figure 1A, with inclusion of MM134+Fulvestrant samples. **(B,D)** Gene set enrichment against GO Biological Processes. Dashed line: adj. p = 0.05. **(C,E)** Mean IgG-subtracted intensity of duplicate IP/MS samples for proteins in indicated ontology groups. Orange bar identifies IP/MS hits putative remodeled by fulvestrant treatment discussed in text.

### Co-regulator activity of MDC1 is linked to specific cellular DNA repair activity

We explored the cross-talk between MDC1 co-regulator activity with ER versus MDC1 DNA repair activity using a single-cell approach to measure DNA repair called “Haircut”. Haircut profiles DNA repair activities using DNA oligo substrates for specific repair enzymes by including oligonucleotide substrates in the single-cell emulsion. Repair enzymes create nicks in the substrates, and sequencing quantifies repair activities by comparing modified and unmodified oligo substrates. Haircut profiles PARP-related single-strand break repair pathways, quantifying the activity of base excision repair and nucleotide excision/incision repair enzymes in a single reaction (in (32) see Figures 1-2 and Supplemental Figure 1, also (33)). We coupled Haircut with targeted transcriptome analysis (10X Genomics Pan-cancer capture panel, n=1,253 genes) to profile DNA repair activity and gene expression in individual cells. We performed scRNAseq + Haircut in MM134 ± estradiol (E2) (n=8,307 cells: –E2, n=4,940; +E2, n=3,367) (**Supplemental File 2** – see access link in methods). We extrapolated ER activity and co-regulator activity (MDC1 vs pioneer factor FOXA1; **Figure 3A**) from the expression of n=259 ER target genes in the capture panel based on our prior RNAseq study (14) (cell cycle genes excluded; **Supplemental File 2**), and examined ER:MDC1 activity versus DNA repair capacities.

**Figure 3.**
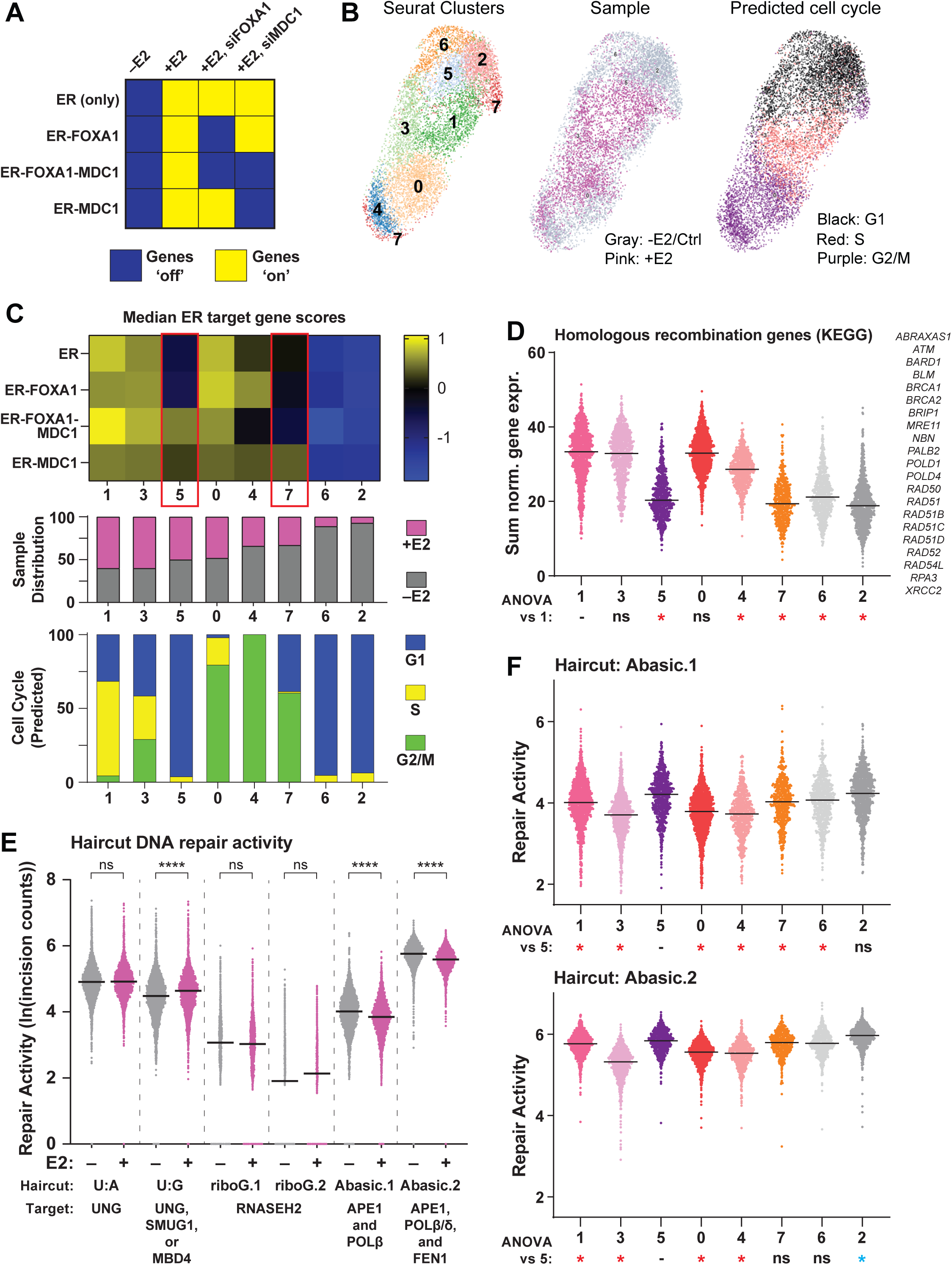
Single cell transcriptome + Haircut data identifies reciprocal MDC1 co-regulator activity versus DNA repair activity. **(A)** Schematic for predicted ER target gene expression based on dependence on MDC1, FOXA1, or neither factor. Based on RNAseq data from Sottnik & Bordeaux et al, 2021. **(B)** UMAP plot showing Seurat clusters (left), sample/treatment (center), and predicted cell cycle state (right). **(C)** Top, ER target gene scores in each of the 8 Seurat clusters. Single cell scores are a z-score of the relative expression of estrogen-induced and estrogen-repressed genes (see Supplemental File 2). Middle, per cluster fraction of cells from the –E2 vs +E2 samples. Bottom, per cluster distribution of predicted cell cycle state. Points represent individual cells. **(D)** Expression of homologous recombination genes (https://www.genome.jp/entry/hsa03440) from targeted capture panel. *, ANOVA adj.p <0.05 versus Cluster 1. Points represent individual cells. **(E)** Haircut assay from hairpin DNA repair substrates that require activity of the indicated enzymes for editing and sequence readout. ****, ANOVA adj.p <0.0001, –E2 vs +E2. Points represent individual cells. **(F)** Repair of the Abasic hairpin yields two outcomes; APE1-mediated incision at an abasic site is followed by processing by either Polβ (short-patch repair, Abasic.1 - top) or Polδ/β and FEN1 (long-patch repair, Abasic.2 - bottom). *, ANOVA adj.p <0.05 versus Cluster 5. Points represent individual cells.

Gene expression via the Pan-cancer capture panel defined 8 clusters (**Figure 3B**) with distinct ER and co-regulator activity (**Figure 3C**; **Supplemental Figure 3A**). Two clusters were primarily –E2 cells with minimal activation of ER target genes (Clusters 6 & 2), while all other clusters had increasing fractions of +E2 cells and showed activation of ER target genes (i.e. “ER-active clusters”). In two ER-active clusters, ER:MDC1 target genes were active while MDC1-independent ER target genes were inactive (**Figure 3C**, clusters 5 & 7; C5/C7). Paralleling this separate activation of MDC1-dependent vs –independent ER target genes, we noted that activation of MDC1-dependent ER targets was effectively independent of cell cycle, whereas MDC1- independent ER targets were activated in S and G2/M versus G1 independent of E2 (**Supplemental Figure 3B**). Specific activation of ER:MDC1 in C5 and C7 and across cell cycle supports that MDC1 has a distinct role in co-regulating ER relative to other factors, i.e., ER:MDC1 activity is separate from ER:FOXA1 and likely is a functional subset of the ER transcriptome.

Notably, C5/7 largely lacked S-phase cells and were enriched for G1 and G2/M cells (as defined by gene expression), respectively. Despite the enrichment for G2/M in C7, both C5/7 expressed markedly low levels of HR-associated genes relative to other ER-active clusters (**Figure 3D**), similar to –E2 clusters 6 & 2. C5/7 were similarly distinct in DNA repair capacity in Haircut analysis. The Haircut “Abasic” reporters primarily measure base excision repair (BER) activity of the enzyme APE1, which cleaves the DNA backbone at abasic sites, and we found that +E2 cells had lower BER/APE1 activity versus –E2 cells (**Figure 3E**). In contrast, C5/7 showed elevated BER/APE1 activity and had the highest such activity among the ER-active clusters (**Figure 3F**). The ER target gene activity and Haircut data taken together link MDC1 co-regulator activity to down-regulated HR activity yet upregulated BER activity, and a potential reliance on non-HR pathways like BER to maintain genomic integrity. These observations parallel the putative HR dysfunction indicated by the MDC1 interactome in ILC cells.

### ILC cells present with dysfunctional initiation and resolution of homologous recombination

The depletion of HR proteins associated with MDC1 in ILC cells, coupled with our prior observation that ILC cells show primarily pan-nuclear γH2AX in response to DNA damage (14), are consistent with DDR or DNA repair dysfunction. To examine the nature of this putative dysfunction further, we induced DNA damage in MM134 (ILC) cells and examined key proteins in DNA damage response initiation and resolution. Etoposide treatment led to DDR protein phosphorylation, including those upstream of MDC1 (H2AX, ATM, MRN complex proteins) and downstream of MDC1 (CHK1/2, 53BP1, DNA-PKcs) (**Figure 4A**), supporting that initiation of DDR is intact in these cells. Notably, phosphorylation of NHEJ-associated proteins (53BP1, DNA-PKcs) was rapid and seen within 4hrs, while RAD51 turnover, associated with HR resolution (34,35), was not apparent in MM134 cells. We next examined induction and resolution of DDR protein phosphorylation in MM134 (ILC) compared to MCF7 (IDC) after etoposide treatment. 53BP1 and DNA-PKcs phosphorylation occurred immediately in MM134 whereas this was delayed in MCF7 (**Figure 4B**), supporting rapid activation of NHEJ in MM134. In parallel, etoposide treatment decreased levels of RAD51 in MCF7, consistent with RAD51 turnover during HR (36), whereas this was not observed in MM134 out to 48hrs post-damage. The lack of RAD51 protein turnover may indicate dysfunctional HR-mediated DNA repair in ILC cells. Based on this, we examined DNA repair foci by immunofluorescence, using RAD51 foci as a marker of HR initiation and resolution (37). We found that ILC cells did form RAD51 foci in response to either ionizing radiation or etoposide, however, foci formation was notably delayed in ILC cells (MM134, 44PE) relative to IDC cells (T47D, HCC1428), in particular in response to ionizing radiation (**Figure 4C; Supplemental Figure 4**). IDC cells formed RAD51 foci within 4 hours of radiation, with marked resolution of RAD51 foci by 24 hours, while ILC cells showed few RAD51 foci until 24 hours post-damage. Of note, these ILC cells have S and G2/M fractions comparable to IDC cell lines (14). Coupled with the lack of RAD51 protein turnover, these data suggest that complete execution and resolution of HR are dysfunctional in ILC cells.

**Figure 4.**
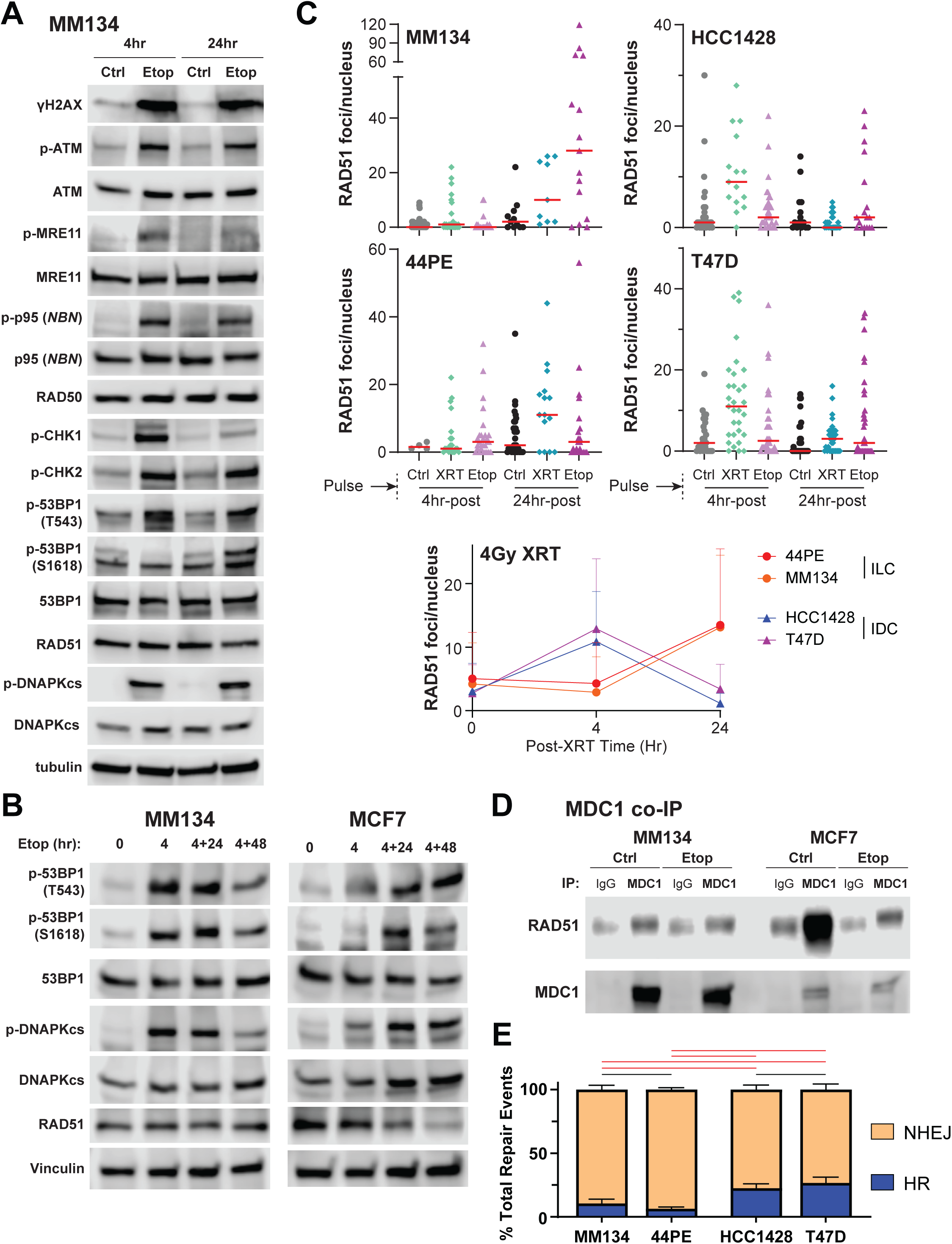
Activation of HR is limited and delayed in MM134 (ILC) versus MCF7 (IDC) cells. **(A)** MM134 cells treated with 10µM etoposide for 4 or 24 hours were analyzed by western blot for canonical interactors of MDC1 (yH2AX, ATM, and MRN) and down-stream mediators of canonical MDC1 signaling (CHK1/2, 53BP1, RAD51, and DNAPKcs). **(B)** Western blot comparison of MM134 (ILC) and MCF7 (IDC) for MDC1 interaction of downstream NHEJ (53BP1 and DNAPKcs) vs HR (Rad51) was performed. Increased turnover of RAD51 in MCF7 is suggestive of HR being used to repair etoposide induced double strand breaks. **(C)** Breast cancer cells were treated with a 4hr pulse of 10μM etoposide, or 4Gy ionizing radiation, and allowed to recover for 4 or 24hr prior to assessing RAD51 foci formation by immunofluorescence. Points represent foci counts in individual nuclei, read line represent median per condition. **(D)** Immunoprecipitation of MDC1 in MM134 and MCF7 cells treated with 10µM etoposide. Increased co-IP of RAD51 is observed in MCF7 but not MM134. **(E)** Traffic Light DNA repair reporter output from breast cancer cells. Bars represent mean ± SD form n=5 biological replicates. Red bar represents ANOVA p<0.05 for indicated comparison.

MDC1 plays a central role in the “decision” between HR and NHEJ, and MDC1:RAD51 association was shown to be critical for HR (36). RAD51 was not detected in our IP/MS studies, so we examined this by co-IP with immunoblotting. We confirmed in MCF7 (IDC) that MDC1 was strongly associated with RAD51, and confirmed the decrease in RAD51 after etoposide treatment (**Figure 4D**). Conversely, we detected minimal MDC1:RAD51 association in MM134 (ILC) and saw no decrease in RAD51 via MDC1 co-IP after DNA damage. Collectively these data are consistent with inefficient or dysfunctional resolution of HR in ILC cells.

To further examine resolution of double-strand break repair via NHEJ versus HR, we utilized the ‘Traffic Light’ I-SceI fluorescent reporter system, which tracks repair pathway choice of NHEJ versus HR at an individual double-strand break per cell (38). We stably integrated the Traffic Light reporter into ILC and IDC cells (see Materials and Methods), and tracked DNA repair after induction of I-SceI by flow cytometry measurement for RFP (NHEJ) versus GFP (HR). NHEJ activity was comparable across ILC and IDC cell lines, however, HR activity was extremely limited in ILC cells; HR accounted for ∼6-10% of repair events in ILC cells, versus ∼25% of events in IDC cells (**Figure 4E, Supplemental Figure 5**; HCC1428 are functionally a *BRCA2*-hypomorph (39)). Taken together, these data support that HR is dysfunctional in ILC cells, and that this dysfunction may not prevent initiation of HR yet ultimately results in ineffective resolution of HR.

### ILC tumors present features of a novel form of DNA repair dysfunction distinct from HR deficiency

Studies to date on HR deficiency related to “BRCAness” do not mechanistically account for the distinct form of dysfunctional, inefficient resolution of HR that presents in our data. As such, we revisited TCGA analyses, including the Pan-Cancer analysis of HR deficiency phenotypes (40), to identify features of ILC that may provide insight into this phenotype and its prevalence across ILC. In the TCGA analyses of Luminal A/ER+ ILC, high-risk ILC with the poorest survival had an elevated DNA Damage Response (DDR) score in reverse phase protein array (RPPA) analyses (4). We examined whether increased DDR protein levels predicted outcomes in ILC versus IDC, and despite limited survival events among ILC in TCGA, high DDR protein levels predicted poorer overall survival in ILC but not IDC (**Figure 5A**). Among these DDR proteins, RAD51 is elevated in ER+ ILC (**Figure 5B-C**) despite significantly lower *RAD51* mRNA levels (**Figure 5D**), suggesting RAD51 is increased post-transcriptionally, e.g. via dysfunctional post-HR turnover. Further, RAD51 protein and mRNA levels are positively correlated in ER+ IDC but no correlation is present in ER+ ILC (**Figure 5E**).

**Figure 5.**
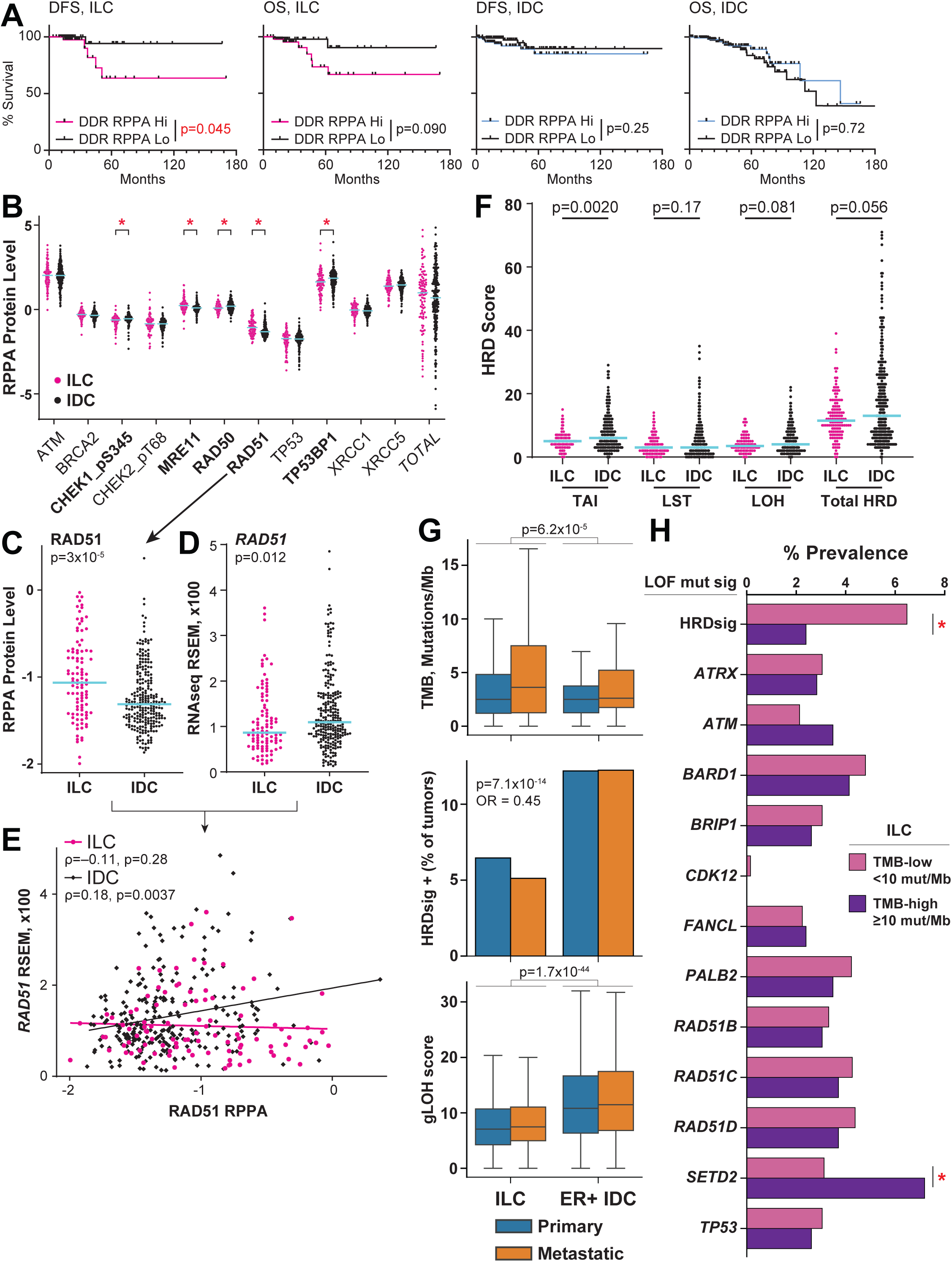
Protein array data are consistent with DNA repair dysfunction in ILC tumors and cell lines. mRNA, protein, and associated clinical data downloaded from cBio Portal (78), TCGA PanCan dataset; HRD feature data from (40). **(A)** Disease-free survival (DFS) and Overall Survival (OS) for Luminal A tumors with available RPPA data. Hi/Lo defined by total RPPA signal sum cutoff of 1. ILC RPPA Hi: DFS n=42, OS n=49; ILC RPPA Lo: DFS n=46, OS n=50; IDC RPPA Hi: DFS n=98, OS n=111; IDC RPPA Lo: DFS n=112, OS n=139. Log-rank test p-value shown. IDC curves are ended at 180mo but no survival events occurred >180mo. **(B)** Normalized RPPA signal shown per target. ILC n=99, IDC n=251. Blue line = median. *, Mann-Whitney with FDR < 5% (adj.p < 0.05). **(C)** RAD51 RPPA data, as in (B). **(D)** *RAD51* mRNA expression. Mann- Whitney T-test p-value shown. **(E)** RPPA vs mRNA data from (C-D); Spearman correlation statistics shown. **(F)** Homologous recombination deficiency (HRD) scores, representative of other trends for DNA damage response signatures observed in Luminal A tumors in (40). ILC n=134; IDC n=315. Blue line = median. Mann- Whitney T-test p-value shown. **(G)** Tumor mutation analyses from Foundation Medicine cohort, details in text. ILC: primary, n=1653; metastatic, n-1535. ER+IDC: primary n=1087; metastatic, n=439. TMB and gLOH scores compared for ILC v IDC (combining primary + metastasis), Mann-Whitney U-test. HRDsig, by prevalence of HRDsig-positive status, compared by Fisher exact test, odds ratio for HRDsig-positive in ILC shown. **(H)** Prevalence of mutational signatures among ILC, per TMB status. TMB-high, n=459; TMB-low, n=2625; not assessable, n=351. *, Fisher exact test adj.p < 0.05.

These observations are consistent with our cell line data and support that resolution of DSB via HR is inefficient or dysfunctional in at least a subset of ILC. However, nearly all genomic markers of HR deficiency were decreased in ILC compared to IDC (40,41), including HR deficiency markers associated with telomeric allelic imbalance (42), large-scale transitions (43), and loss-of-heterozygosity events (44) (**Figure 5F**). Collectively, the absence of genomic scarring associated with overt HR deficiency in ILC tumors is consistent with our mechanistic data.

Previous analysis of the Foundation Medicine database (FoundationCORE®) in 2019 (then with n=516 ILC) showed that ILC carry an increased tumor mutational burden (TMB) versus ER+ IDC, in particular for metastatic ILC. We reanalyzed TMB data from FoundationCORE (updated to include n=3,435 ILC and n=19,429 IDC) to determine whether the elevated TMB in ILC is associated with mutational signatures of loss-of-function of specific DDR genes to provide insight into the DNA repair dysfunction phenotype. Signatures associated with biallelic loss of function of DDR genes (n=12) were developed using an extreme gradient boosting model (XGBoost, see (45) and Methods). Of note, confirmed ER status is not typically available for this cohort, but since ILC are ∼95% ER+, we focused on IDC with known ER+ status (n=1,645). In the updated cohort, TMB is increased in ILC versus ER+ IDC (**Figure 5G**). Despite this increased TMB, as observed in TCGA, markers of HR deficiency including genome-wide LOH events and scar-based HRDsig (46) were decreased in ILC vs ER+ IDC (**Figure 5G**). Similarly, loss-of-function signatures for nearly all DDR genes examined were less common in ILC than IDC or ER+ IDC, with the exception of *SETD2*, which was present in 3.8% of ILC versus 1.2% of IDC or 1.4% of ER+ IDC (ILC vs ER+ IDC chi test p=2.6×10^-6^; **Supplemental Figure 6A**). In both ILC and ER+ IDC, tumors positive for HRDsig or loss-of-function signatures for *BARD1*, *BRIP1*, *PALB2*, *RAD51B/C/D*, and *TP53* had modest but significant increases in TMB relative to signature-negative tumors. The *ATM* signature was linked to increased TMB in ILC but not ER+ IDC, and the *SETD2* signature showed a trend for increased TMB in ILC but decreased TMB in ER+ IDC (**Supplemental Figure 6B**). However, these increases overall did not lead to clinically-relevant high TMB (i.e. TMB-high, ≥10 mutations/Mb). Comparing TMB-high ILC (15% of tumors) vs TMB-low ILC (85%), we noted that HRDsig- positive tumors were significantly depleted among TMB-high ILC, and nearly all other signatures were modestly depleted among TMB-high ILC (**Figure 5H**). The exceptions were a modest enrichment of *ATM* signature positive tumors and a significant, >2-fold enrichment of *SETD2* signature positive tumors among TMB-high ILC (**Figure 5H**). As reported previously, nearly all TMB-high tumors showed an APOBEC-driven mutational signature (**Supplemental Figure 6C**). These data show that ILC do not present with typical genomic features of HR deficiency and that TMB in ILC is independent of overt HR deficiency, together supporting that the mechanisms of DDR/HR dysfunction in ILC are distinct among breast cancers.

### ILC cells are sensitive to PARP inhibitor talazoparib in vitro and in vivo

From the TCGA Pan-Cancer HR deficiency study, the only signature elevated in ILC is PARPi7 score (**Figure 6A**). PARPi7 is a normalized gene expression score predictive of sensitivity to PARP inhibitors in I-SPY trials ((47); high *CHEK2* and *MAPKAPK2*, with low *BRCA1*, *MRE11*, *NBN*, *TDG*, and *XPA*, drive increased PARPi7). High PARPi7 score predicted pathologic complete response in high-risk, ER+ breast cancer in response to PARPi veliparib + carboplatin (48) or PARPi olaparib + durvalumab (49). Notably, the increased PARPi7 in Luminal A ILC (median = 0.158) is likely functionally significant, as cutoffs of 0.0372 and 0.174 separated PARPi-sensitive versus -resistant tumors using Affymetrix and Agilent data, respectively (47,48).

**Figure 6.**
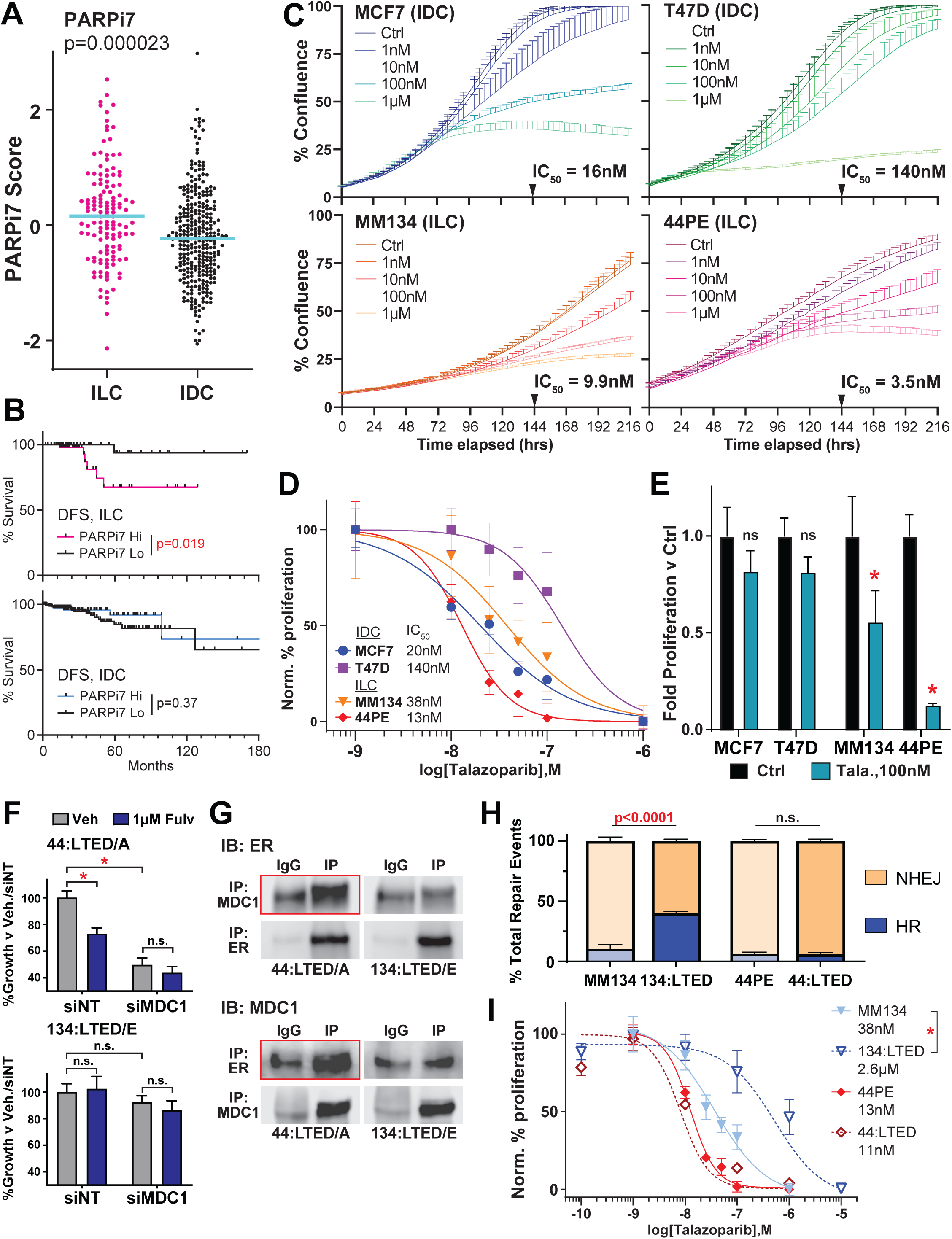
ILC cells *in vitro* are sensitive to PARP inhibitor talazoparib. **(A)** PARPi7 score from TCGA PanCancer analysis (40). ILC n=134; IDC n=315. Blue line = median. Mann-Whitney T-test p-value shown. **(B)** DFS from samples with PARPi7 score and survival data. ILC PARPi7 Hi, n=57; ILC PARPi7 Lo, n=62; IDC PARPi7 Hi, n=82; IDC PARPi7 Lo, n=188. Log-rank test p-value shown. IDC curves are ended at 180mo but no survival events occurred >180mo. **(C)** Representative growth curves via confluence in live cell imaging (Incucyte) for ILC vs IDC cells with increasing concentrations of talazoparib. IC_50_ calculated using percent confluence at 144hr post-treatment. Line represents mean confluence of 6 replicates ± SD. **(D)** Proliferation at 7d (168hr) as assessed by dsDNA quantification, shown as normalized % proliferation vs max/min. Points represent mean of 6 replicates ± SD. **(E)** Proliferation at 7d post drug washout (14d after initial treatment) by dsDNA quantification. Bars represent mean of 6 replicates ± SD. *, ANOVA, Ctrl v Tala p<0.05. **(F)** Proliferation at 7d as in (D-E) after siRNA transfection or indicated treatment. /A and /E designations indicate representative LTED sub-lines used in studies herein, from (51). **(G)** Reciprocal co-immunoprecipitation of MDC1 versus ERα (ER). Red boxes highlight ‘target’ co-IPs confirming ER:MDC1 association in 44:LTED/A, which is not enriched versus IgG control in 134:LTED/E. **(H)** Traffic Light reporter as in Figure 4E, data for parental cells replicated from Figure 4E for clarity. ANOVA p-value shown. **(I)** Proliferation at 7d with increasing talazoparib concentration, as above. Dose-response curves for parental cells from Figure 6B shown for clarity. *, IC_50_ comparison p<0.05.

Absent a trial-defined cutoff for RNAseq data, at a more conservative cutoff of 0.2, 49% of Luminal A ILC vs 28% of Luminal A IDC are PARPi7-high (Χ^2^ test, p=0.000018); this cutoff identified poor disease-free survival in ILC, but not IDC (**Figure 6B**). Despite the disconnect between PARPi7 score and other markers of genomic scarring, this observation suggests that HR dysfunction in ILC cells causes sensitivity to PARPi. We examined whether ILC cells are sensitive to PARP inhibition, focusing on FDA-approved agent talazoparib, which more potently traps PARP1/2 on DNA relative to olaparib (50). Via live cell Incucyte imaging we found that ILC cell lines MM134 and 44PE are growth-suppressed by low-nanomolar concentrations of talazoparib (**Figure 6C**), similar to MCF-7 but ∼10-fold more sensitive than T47D cells. Parallel experiments measuring cell proliferation by dsDNA quantification confirmed low-nanomolar talazoparib sensitivity in ILC cells (**Figure 6D**, **Supplemental Figure 7A**). Notably, both confluence and dsDNA quantification time-courses suggested that a differential impact of talazoparib on ILC cells manifested at 5-6 days post-treatment; IDC cells appeared viable despite growth suppression, but ILC cells appeared non-viable after talazoparib treatment (**Supplemental Figure 7A-B**). To examine post-treatment cell viability, we washed out drug and re-plated cells after the 7 day treatment, and assessed outgrowth. ILC cell lines MM134 and 44PE, but not IDC lines MCF7 and T47D, remained significantly growth-inhibited after drug washout (**Figure 6E**), suggesting that PARPi treatment had a long-term impact on cell viability specifically in ILC cells.

Our data suggests that PARPi sensitivity in ILC is linked to ER:MDC1 activity and HR capacity, which we were able to examine using our long-term estrogen deprived variants (LTED, modeling aromatase inhibitor resistance) of MM134 (134:LTED) and 44PE (44:LTED) (51,52). We previously showed that though these models are ER+, 134:LTED are ER-independent while 44:LTED are ER-dependent. Accordingly, 134:LTED are not growth-inhibited by fulvestrant or MDC1 knockdown, while 44:LTED remain sensitive to both fulvestrant and MDC1 knockdown (**Figure 6F**), and the ER:MDC1 interaction is lost in 134:LTED but maintained in 44:LTED, respectively (**Figure 6G**). These observations suggest MDC1 is maintained as a transcriptional partner of ER in 44:LTED, but is de-coupled from ER in 134:LTED and thus may re-engage in DDR activity. Using the Traffic Light reporter, we found that 44:LTED perform limited HR as seen in parental 44PE, but HR activity is strongly increased in 134:LTED, consistent with a restoration of HR activity (**Figure 6H**). Restoration of HR activity was accompanied by a decrease in sensitivity to talazoparib specifically in 134:LTED, which were >50-fold less sensitive to the PARPi versus parental MM134 (**Figure 6I**). Together these data support that differential activity of MDC1 in DDR underpins responsiveness to PARPi in ILC cells.

We next examined single-agent talazoparib efficacy *in vivo* using mammary fat pad xenograft tumors of MM134 cells in female immunocompromised mice (NSG, NOD.Cg-*Prkdc^scid^ Il2rg^tm1Wjl^*/SzJ). 20 mice were injected with 5×10^6^ MM134 cells in matrigel bilaterally at the #4 mammary fat pad, and mice were supplemented with 30μM estradiol in drinking water to support tumor growth. Treatments were initiated at day 27 post-tumor challenge with >25% of tumors >100mm^3^. n=5 mice per arm received vehicle (vehicle gavage, see Methods), talazoparib (0.33mg/kg; 5d/w p.o.) for 3 weeks, talazoparib for 6 weeks, or fulvestrant (5mg/kg s.c. weekly) for 6 weeks; tumors were tracked until all vehicle control mice reached humane study endpoints (day 79). Of note, the 6-week talazoparib arm was lost at day 72 due to a cage flood and data are censored and excluded from analysis. In this study, single agent talazoparib (Tala) treatment significantly suppressed tumor growth versus control (ANOVA Vehicle vs 3wk Tala, p<0.0001), and outperformed fulvestrant in reducing tumor growth (ANOVA Fulvestrant vs 3wk Tala, p=0.0035) (**Supplemental Figure 8A**). Fulvestrant provided an initial growth suppression, but several tumors continued to grow on active treatment. Conversely, though talazoparib treated tumors were larger than at study initiation (3wk Tala: ∼2.9-fold increase at endpoint, paired t-test p=0.002), growth inhibition was durable and tumors did not significantly increase in volume between treatment completion and study endpoint (day 48 vs day 79, paired t-test p=0.11).

We next tested the efficacy of talazoparib alone or in combination with endocrine therapy, via fulvestrant or estrogen withdrawal (to model aromatase inhibitors, AI) against MM134 and 44PE xenografts. Treatment was maintained for 6 weeks and then tumor outgrowth was tracked after cessation. Tala alone suppressed tumor growth vs control in MM134 (ANOVA p=0.0016) but not 44PE; adding fulvestrant to Tala showed no additive effect for either cell line (**Supplemental Figure 8B-C**). However, combined AI+Tala was highly effective in both models, with AI+Tala superior to either single treatment against tumor growth (**Figure 7A-D**), and caused a durable slowing of tumor outgrowth after treatment cessation (**Figure 7A-B**). Both cell line xenografts were growth inhibited by AI (estrogen withdrawal), however, AI+Tala caused a greater decrease in tumor volume at endpoint for both models (**Figure 7C-D**). Estrogen withdrawal (AI)-only caused a modest decrease in tumor volume in 44PE with 1/10 tumors increasing in size by treatment completion (25±37% decreased volume, paired ANOVA p=0.06), while AI+Tala was more effective (56±27% decreased volume, paired ANOVA p=0.0006). These data support that ILC are sensitive to PARPi despite lacking HR-deficiency.

**Figure 7.**
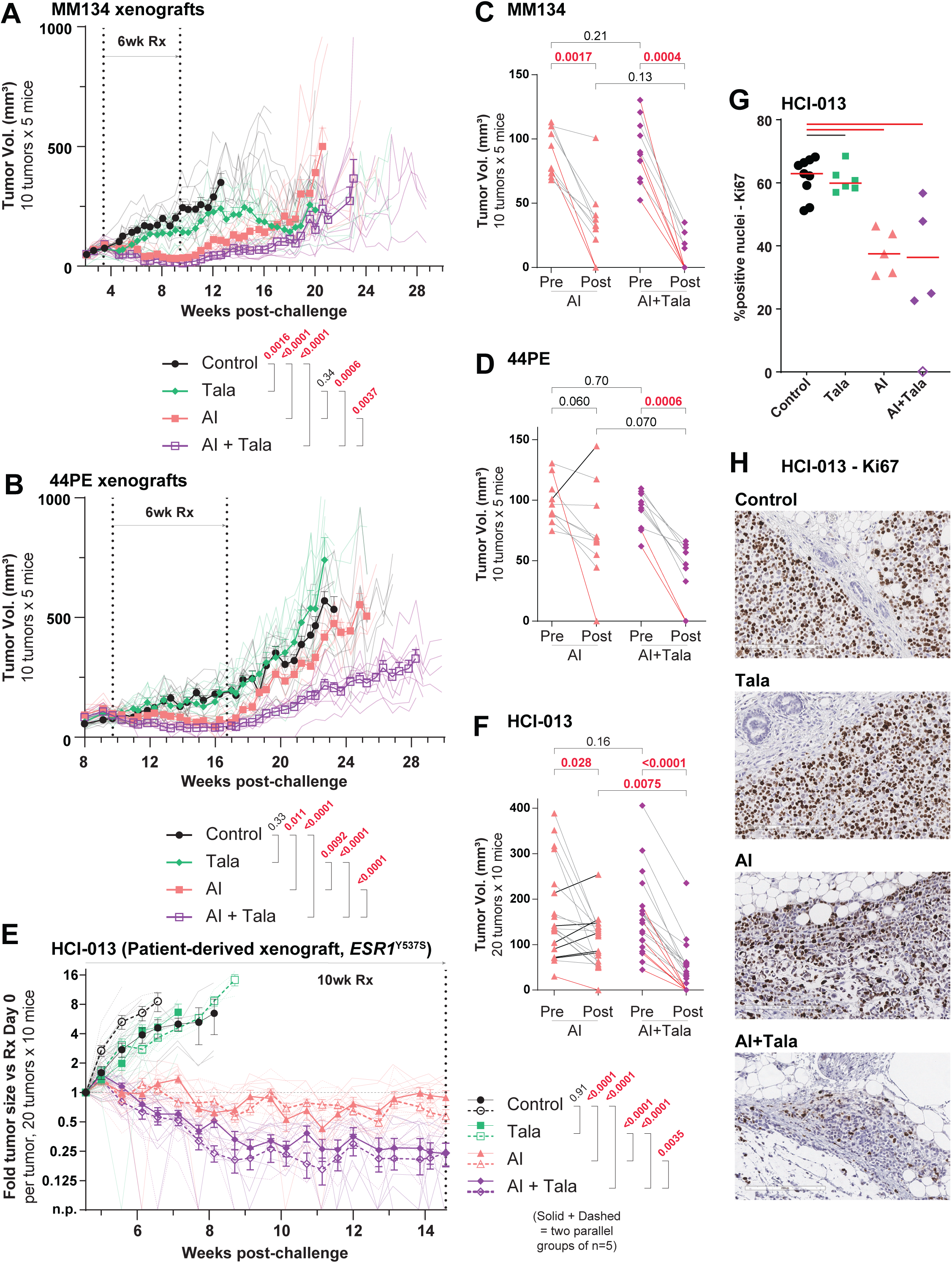
Talazoparib causes sustained growth suppression in ILC xenograft tumors. (A-B) Bold lines/symbols show mean tumor size ± SEM; individual tumor size shown as matching faded lines. Bold lines ended at first tumor size human endpoint reached per arm. Two-way ANOVA treatment effect p-values shown. **(C-D**) Tumor volume at treatment initiation vs completion for studies in (A-B). Red lines = tumor non-palpable at treatment completion; Black lines = increased tumor volume at treatment completion vs initiation. Paired ANOVA p-values shown. **(E)** Tumor growth shown as fold-change in tumor size versus treatment initiation.

Testing PARPi combinations *in vivo* against anti-estrogen-resistant ILC models is critical, however, our LTED variant have shown limited engraftment as mammary fat pad xenografts in contrast to the parental cell lines (not shown). To examine PARPi efficacy against anti-estrogen-resistant ILC, we utilized patient-derived xenograft (PDX) model HCI-013, which was derived after that patient had received letrozole (AI) + leuprolide, tamoxifen, and exemestane (AI), followed by several chemotherapy regimens; HCI-013 also carries the *ESR1*^Y537S^ mutation (53,54). After 10 weeks treatment with Tala ± AI (estrogen withdrawal), AI+Tala caused the strongest suppression of PDX growth (**Figure 7E-F**). AI alone did limit tumor growth versus control (**Figure 7E**), but 30% of tumors (6/20) had increased in size over the treatment period (**Figure 7F**). No progression was observed with AI+Tala and 30% of tumors (6/20) were non-palpable at treatment completion, while AI+Tala-treated tumors were significantly smaller than AI alone at endpoint (**Figure 7F**) (AI: 27±42% decreased volume, versus AI+Tala: 76±20% decreased volume; post-treatment volume, AI: 106±55mm^3^ versus AI+Tala: 46±55mm^3^, ANOVA p<0.0001). Small lesions in AI+Tala treated mice showed limited cellularity in the tumor bed (**Supplemental Figure 9**). Immunohistochemistry for tumor proliferative marker Ki67 at treatment completion showed that AI or AI+Tala suppressed cell proliferation versus Control (**Figure 7G-H; Supplemental Figure 9**). Proliferation in AI+Tala was not significantly different than AI alone, but residual lesions showed less tumor cellularity (**Supplemental Figure 9**). Together these data support that ILC have an unexpected sensitivity to PARPi that can be exploited using aromatase inhibition combined with talazoparib.

## DISCUSSION

Patients diagnosed with ILC face distinct treatment and clinical management challenges, in part related to uniquely poor long-term outcomes, controversy around the efficacy of anti-estrogens, and a lack of treatment strategies based on ILC biology. Toward better understanding estrogen and anti-estrogen response in ILC, we previously identified that the DNA repair protein MDC1 has novel ER co-regulator activity in ILC cells which mediates estrogen response and anti-estrogen resistance, yet is associated with putative DNA repair dysfunction (14). In the present study, we profiled the MDC1 interactome to characterize MDC1 DNA repair versus transcriptional regulator functions, in IDC versus ILC cells. MDC1 associations with homologous recombination proteins were depleted in ILC cells, mirroring that observed in *BRCA2*-mutant IDC cells. Using single-cell transcriptomics paired with DNA repair activity measurement with the ‘Haircut’ assay, we found that ER:MDC1 gene regulation activity was associated with an upregulation of PARP-mediated repair pathways, specifically base excision repair. Functional studies of DDR initiation, propagation, and resolution support that ILC cells have a distinct form of DNA repair dysfunction, and we tested whether this dysfunction could be exploited using FDA-approved PARPi talazoparib. Both *in vitro* and *in vivo*, ILC cells were sensitive to talazoparib, which caused durable suppression of cell/tumor growth after treatment cessation. Understanding how MDC1 underpins ILC-specific ER functions and DNA repair dysfunction offers a path toward ILC precision therapies based on identifying tumors where MDC1 activity indicates PARPi sensitivity.

ILC are not associated with typical features of DNA repair deficiency or sensitivity to DNA damage – in addition to the limited genomic scars of overt HR deficiency we noted in TCGA and Foundation Medicine data and depletion of associated germline mutations in ILC (20–22), ILC generally have limited response to chemotherapy (2,55). However, our data taken together with observations in the literature suggest that ILC present with DNA repair dysfunction that does not manifest as HR deficiency (HRD) as canonically understood. HR dysfunction in ILC may be linked to the elevated DNA damage response signature (i.e. DDR protein levels) reported in TCGA analyses in high-risk ILC, which supports that DDR plays a distinct role in ILC etiology. The mechanistic significance of these elevated DDR protein levels in high-risk ILC is unknown, but the lack of RAD51 turnover we observed suggests a link to inefficient or dysfunctional resolution of HR. This more subtle dysfunction may not be captured in current HRD signatures, which are largely trained off of HR-deficient (*BRCA1/2*-mutant) tumors. In that context, mutations in other HR genes are often linked to high HRD scores in a tissue type-dependent manner (56), so ILC may also have unique associations with high HRD scores.

Similarly, we observed that ILC cells formed RAD51 foci upon damage (which would otherwise suggest ILC cells are HR-proficient (37)) despite showing little HR-mediated repair in I-SceI (Traffic Light) reporter assays. Identifying the point of dysfunction in HR is an important future direction. The restoration of the HR-associated Bold/faded lines as above; two-way ANOVA includes all 20 tumors per treatment arm together. n.p. = non-palpable at measurement timepoint (volume = 0 for ANOVA). **(F)** Change in tumor volume for HCI-013 as in (C-D). **(G)** Ki67 quantification from tumors in (E,F); control and tala-treated tumors were collected asynchronously at humane tumor size endpoints. AI and AI+Tala-treated tumors represent the dashed-line cohort from (E); open symbol in AI+Tala was 0% Ki67+ but had <10 evaluable tumor cells in available sections, and is excluded from statistical comparisons. Non-parametric ANOVA shown; black line, p>0.05; red lines, p<0.05. **(H)** Representative Ki67 immunohistochemistry from tumors in (E,F,G), scale bar = 200μm; also see Supplemental Figure 9.

MDC1 interactome upon fulvestrant treatment, and restoration of HR activity in 134:LTED cells, suggests that the HR dysfunction is dynamic or transient, via active regulation of MDC1 or associated factors. Since restoration of HR in 134:LTED was associated with talazoparib resistance in those cells, understanding HR dysfunction and regulation is critical toward understanding PARPi sensitivity in ILC.

Defining the connection between HR dysfunction and tumor mutational burden (TMB) in ILC may also yield important insight into DNA repair capacity and PARPi sensitivity in ILC. The increased TMB in ILC is not likely driven by a subset of tumors with overt HR deficiency, since scarring (HRDsig) was significantly depleted among TMB-high ILC, as were most DDR loss-of-function mutational signatures. The enrichment of *ATM* and *SETD2* loss-of-function signatures in TMB-high ILC may point to mechanisms for HR dysfunction in ILC, but germline *ATM* mutations are in <1% of ILC and not enriched vs IDC (22); *SETD2* mutations are uncommon but may be enriched in metastatic ILC (∼3%) vs metastatic IDC (∼1%) (57). An enrichment in APOBEC mutational signatures in ILC versus IDC tumors may also underpin high TMB. Sokol et al linked the elevated TMB observed in metastatic ILC to APOBEC, as 88% of TMB-high metastatic ILC had a dominant APOBEC signature (24), which we confirmed in the extended cohort analyses herein. APOBEC signatures are enriched in ∼16-38% of primary ILC (vs 8-16% of primary IDC/NST) and 22-51% of metastatic ILC (vs ∼10- 28% of metastatic IDC/NST) and are associated with increased TMB relative to APOBEC-negative tumors (57–59). APOBEC lesions are repaired by PARP-driven BER pathways, and high APOBEC activity has been associated with PARPi-sensitivity (60,61); high APOBEC activity combined with HR dysfunction may impart PARPi sensitivity in ILC without otherwise driving broad HRD-related scarring.

Though PARP inhibitors including talazoparib have been widely tested in breast cancer, their clinical efficacy for ILC, especially beyond *BRCA1/2*-mutant tumors, is unknown. Neither the phase III olaparib trial OlympiAD nor the two largest breast cancer trials of talazoparib to date (EMBRACA (62) and ABRAZO (63)) reported tumor histology. A pilot neoadjuvant talazoparib study (64) included one germline *BRCA2*-mutant patient with ER+ ILC which had a pathologic complete response to talazoparib, but the extended version of the trial, NEOTALA, did not report tumor histology (65). Defining biomarkers for DNA repair activity and/or MDC1 repair vs genomic activity in ILC is critical to better understand the scope of DNA repair dysfunction in ILC and to design effective trials for talazoparib in ILC.

Though *TP53* mutations are uncommon in ILC, the cell line models used here (with the exception of MCF7) do carry *TP53* mutations, which is typical of cancer cell lines including ILC (66–68). MM134 and 44PE do mirror ‘Proliferative’ ILC regarding the elevated DNA damage response protein signature (69), yet few other models of ER+/HER2-negative ILC exist. Recent multi-omic profiling of putative ILC lines supports that among 5 luminal-like ILC cell lines, MM134 and 44PE are suitable models of ER+/luminal ILC, while BCK4 and MDA MB 330 are HER2-positive (via mutation and amplification, respectively), and CAMA1 has little transcriptional similarity to ILC tumors (68). PDX of ER+ ILC are similarly limited; HCI-013 is the only ER+ classic ILC PDX (i.e. not mixed ILC/NST) used in drug treatment and endocrine-response studies in the literature to date (53,66). Functional studies of DDR/HR and PARPi response in additional models as ILC become increasingly represented will be important to dissect DDR capacity and HR-related biomarkers.

The increasing urgency to develop treatment strategies centered on a differential diagnosis of breast cancer of no special type (i.e. IDC) versus ILC must be addressed through deeper mechanistic understanding of ILC biology. In contrast to the lack of histology information in most trials, recent clinical studies have enriched for or are specifically enrolling patients with ILC (e.g. ROSALINE (70), GELATO (71), NCT02206984). New prognostic signatures developed based on ILC tumors have been reported (72), and pre-clinical studies focusing on ILC models are beginning to identify and define ILC-specific biology toward ILC-tailored treatments (14,52,73–75). Our data advance this paradigm of targeting ILC biology, and show that ILC-specific association between ER and MDC1 creates a new context for ER and MDC1 function in ILC, yet creates a putative synthetic vulnerability via DNA repair dysfunction that may be exploitable with PARPi.

## MATERIALS AND METHODS

### Cell culture

MDA MB 134VI (MM134; ATCC HTB-23) and SUM44PE (44PE; Asterand/BioIVT) were maintained as described (53). HCC1428 (ATCC CRL-2327), MCF7 (Rae Lab, U. Michigan) and T47D (Sartorius Lab, U. Colorado) were maintained as described (14). 134:LTED (subline 134:LTED/E) and 44:LTED (subline 44:LTED/A) were maintained in estrogen-depleted conditions as described (51). 293FT cells (Invitrogen) were maintained according to manufacturer directions. Cells were incubated at 37°C in 5% CO_2_. All lines were regularly confirmed mycoplasma negative (MycoAlert, Lonza), and authenticated by STR profiling at the U. Colorado Anschutz Tissue Culture Core. Estradiol (E2) was from Sigma; ICI 182780 (fulvestrant; fulv) was from Tocris Biosciences. Etoposide and talazoparib (BMN-673) were from Cayman Chemical. Small molecules were dissolved in ethanol or DMSO, and vehicle treatments use 0.01-0.1% EtOH or DMSO *in vitro*.

### MDC1 co-immunoprecipitation and mass spectrometry

Immuno-precipitation (IP) was performed as previously described (14). Briefly, nuclear extract (NE) was prepared from successive hypotonic/hypertonic washes and lysing, then utilized for immunoprecipitation. 10μg antibody was used with NE for immunoprecipitation. Jackson ImmunoResearch Chromapure Mouse IgG was used for control. α-MDC1 (Sigma M2444) was used for MDC1 IP (**Supplemental Figure 10**). NE was incubated with antibody overnight at 4°C with rotation, then extracted with magnetic A/G beads for 4hr at 4°C with rotation. For IP/mass spectrometry, bead-complexes were submitted intact to the Mass Spectrometry Core Facility at the University of Colorado – Boulder for sample preparation and analysis, using an Orbitrap nanoLC-MS. For IP followed by immunoblot, protein complexes were eluted by re-suspending beads in Laemmli-based buffer.

MDC1-IP/MS iBAQ values were used for comparison. IgG iBAQ values were subtracted as background from MDC1-IP values for all proteins. Only proteins enriched (i.e. background subtracted iBAQ > 0) were included in subsequent analyses. Genes present in GO genesets GO:0006397, GO:0000398, GO:0000377, GO:0006364, and GO:0042254 (related to RNA metabolism, see Supplemental File 1) were excluded from analysis as indicated in the text. MDC1 co-immunoprecipitated genes in 293FT (n=1666) were compared to Salifou et al (27) for validation of our IP/MS method. Enriched gene lists for each cell line were analyzed using EnrichR (76) for gene set enrichment analysis. Hierarchical clustering of gene sets was performed using Morpheus (https://software.broadinstitute.org/morpheus).

### Single cell transcriptomics and Haircut analyses

MM134 cells were hormone-deprived in medium supplemented with charcoal-stripped FBS as previously described (77), then treated with vehicle (0.01% ethanol) or 100pM estradiol for 24hrs. After treatment, single cell suspensions were prepared according to 10X Genomics recommendations, and as described (32,33). Whole transcriptome and Haircut libraries were prepared as described (32,33), and separate targeted capture libraries were prepared according to the manufacturer’s instructions (https://www.10xgenomics.com/products/targeted-gene-expression). Supplemental File 2 includes workflow information and normalized gene expression data for the targeted capture panel and Haircut probes, with single-cell annotation for clustering, cell cycle prediction, and ER target genes scores, and is available at: https://osf.io/rw2n6. Raw data will be made available upon submission for publication, or upon request.

### Immunoblotting

Cells were seeded in 12-well plates for ∼80% confluence (MM134: 800k/well; MCF7: 320k/well) and treated as indicated prior to lysis with RIPA buffer (Thermo Pierce). Primary antibodies were diluted to manufacturer recommended concentrations, in TBS + 0.05% Tween-20. Membranes were imaged on LiCor C-DiGit blot scanner. Primary antibodies for immunoblots were: phospho-Histone H2A.X (Ser139)(20E3) (Cell Signaling Technology #9718); ATM (D2E2) (Cell Signaling Technology #2873); Phospho-ATM (Ser1981)(D6H9)(Cell Signaling Technology #5883); MRE11 (31H4)(Cell Signaling Technology #4847); phosph-MRE11 (Ser676)(Cell Signaling Technology #4859); p95/NBS1 (NBN) (D6J5I)(Cell Signaling Technology #14956); phospho-p95/NBS1 (NBN)(Ser343)(Cell Signaling Technology #3001); Rad50 (Cell Signaling Technology #3427); phospho-CHK1 (Ser345)(133D3)(Cell Signaling Technology #2348); phospho- CHK2 (Thr68)(C13C1)(Cell Signaling Technology #2197); 53BP1 (E7N5D) XP (Cell Signaling Technology #88439); phospho-53BP1 (Ser1618)(D4H11)(Cell Signaling Technology #6209); phospho-53BP1 (Thr543)(Cell Signaling Technology #3428); RAD51 (D4B10)(Cell Signaling Technology #8875); DNA-PKcs (3H6)(Cell Signaling Technology #12311); phospho-DNA-PKcs (Ser2056)(E9J4G)(Cell Signaling Technology #68716); α/β-Tubulin (Cell Signaling Technology #2148); Vinculin (E1E9V) XP (Cell Signaling Technology #13901); MDC1 (MDC1-50)(Sigma-Aldrich M2444).

### DNA damage response and HRD scores

DNA repair protein levels, and associated clinical and tumor data, from the TCGA PanCancer dataset were downloaded from cBio portal (78); data were most recently downloaded for verification in October 2024. Scores for genomic markers of HRD including PARPi7 score and associated scarring phenotypes (40) were downloaded from the Genomic Data Commons (https://gdc.cancer.gov/about-data/publications/PanCan-DDR-2018) and a combined clinical/genomic TCGA dataset from the Gerke Lab (https://github.com/GerkeLab/TCGAhrd).

### Foundation Medicine Data Set (FoundationCORE)

Tumor samples were sequenced by hybrid capture–based CGP in a Clinical Laboratory Improvement Amendments– certified/CAP-accredited laboratory (Foundation Medicine, Cambridge, MA) as part of routine clinical care. Sequencing of tissue biopsy was performed on ≥50 ng DNA extracted from 40µm of formalin-fixed paraffin-embedded (FFPE) sections to create adapter sequencing libraries before hybrid capture and sample-multiplexed sequencing for at least 295 genes. Results were analyzed for base substitutions, short indels, CN alterations, and rearrangements. Approval for this study, including a waiver of informed consent and a HIPAA waiver of authorization, was obtained from the Western Institutional Review Board (Protocol No. 20152817). The work conforms to the principles of the Helsinki Declaration.

### XGBoost models to predict biallelic gene loss-of-function features

For each gene of interest, a copy number feature based model was trained to predict biallelic loss. The FoundationCORE data of 434,454 samples was divided into three groups: samples with biallelic alterations in the gene of interest (positive), samples with no alterations in the gene of interest (negative), and samples with alterations that could not be determined to be biallelic (ambiguous). The positive and negative cohorts were divided into train/test split (70/30), and the ambiguous samples were held out. Copy number features for each sample were determined as described (45,79) and used as input to construct the extreme gradient boosting (XGB) model using the caret package in R. Performance was evaluated by ROC-AUC. In total, 18 individual gene models were trained.

Models were considered biologically meaningful if the AUC exceeded 0.7. The ideal cut-off for predicting gene loss was defined as the maximum threshold where sensitivity was at least 70% and the specificity was 95%. Co-occurrence (fisher exact, FDR corrected) for biallelic alterations in the cohort was also tested to confirm that biallelic loss was significantly enriched (p<0.05) in the predicted positive cohort. Six gene models were excluded by AUC, and a further two (*ATRX* and *FANCL*) were underpowered for co-occurrence and were also excluded from further analysis.

### Immunofluorescence

Cells were plated [MCF7 (2×10^4^ cells per well), 44PE (1×10^4^ cells per well), and MM134/44PE (1×10^5^ cells per well)] in 8-well chamber slides (Nunc Tek Chamber Slide System; Thermo Scientific). 24 hours after plating, cells were treated with 0.01% DMSO, 10µM etoposide, or 4Gy of XRT (Multirad350 irradiator). At time points noted, media was aspirated, and cells fixed with 4% methanol free paraformaldehyde (PFA; Electron Microscopy services) for 10 minutes at room temperature. Slides were permeabilized with 0.2% Triton X-100 in PBS. Anti-RAD51 antibody (Abcam, ab176458) was diluted 1:500 in 3% BSA and incubated for 2 hours at room temperature (RT). Goat anti-rabbit Alexa-Fluor 488 conjugated secondary antibody (ThermoScientific; A32731) was diluted 1:1000 in 3% BSA, and incubated on slides for 1 hour at RT. Slides were mounted with SlowFade™ Diamond Antifade with DAPI (ThermoFisher Scientific) for flurescence imaging. CellProfiler software (80) was used for foci analysis. A DAPI mask was used to isolate nuclei as regions of interest for defining RAD51 foci.

### Traffic Light Reporter Assay

The Traffic Light Assay (38) was performed as described, but the Traffic Light reporter was stably integrated into MM134, 44PE, HCC1428, T47D, 134:LTED, and 44:LTED cells. I-SceI was transiently expressed to induce the double-strand break in the reporter via transient lentiviral transduction using an integrase-dead lentivirus (see below). 72 hours after transduction and I-SceI expression, cells were trypsinized, washed, and diluted in FACS buffer (PBS + 1mM EDTA + 25mM HEPES + 1% BSA) for flow cytometry.

Repair events were counted via red vs green fluorescence compared to cells without transient I-SceI expression, on a Beckman Coulter Gallios561 cytometer (Indianapolis, IN, USA) at the U. Colorado Cancer Center Flow Cytometry Shared Resource. Data were analyzed using Kaluza Analysis Software v2.1 (Beckman Coulter).

Traffic Light Reporter (pCVL Traffic Light Reporter 2.1, Addgene #31483) and companion I-SceI vector (Addgene #31476) were a gift from Andrew Scharenberg. For Traffic Light reporter transduction, lentivirus was constructed in 293FT cells via co-transfection with psPAX2 (Addgene #12260, a gift from Didier Trono) and pMD2.G (Addgene #12259, a gift from Didier Trono). For transient I-SceI vector transduction, psPAX2 was replaced with psPAX2-D64V (Addgene #63586, a gift from David Rawlings & Andrew Scharenberg).

### Cell proliferation and viability

Total double-stranded DNA was quantified by Hoechst 33258 fluorescence as described (81), as a surrogate for cell number. Live cell analysis was performed using an S3 Incucyte (Sartorius, Bohemia, NY, USA) with the U. Colorado Cancer Center Cell Technologies Shared Resource. MCF7 (2×10^3^ cells per well), T47D (3×10^3^ cells per well), MM134 (2×10^4^ cells per well), and 44PE (1.5×10^4^ cells per well) were plated in a 96-well format. 24 hours after plating, cells were treated with the noted concentrations of talazoparib and placed in the S3 Incucyte for 216 hours. For analysis, a confluence phase mask was used to determine relative confluency. Alternatively, cells were pre-treated with talazoparib for 7 days prior to re-plating in the 96-well format, and subsequent analysis.

### Animals and xenograft treatment studies

All animal studies were performed in in AALAC-approved facility with approval of the University of Colorado – Anschutz Medical Center Institutional Animal Care and Use Committee (IACUC). 6-8 week female NSG (NOD.Cg-*Prkdc^scid^ Il2rg^tm1Wjl^*/SzJ; Jackson Laboratory strain 005557) mice were used for all experiments. Five mice per group were used for all *in vivo* experiments.

One week prior to tumor challenge, mice were initiated on 30µM 17β-estradiol (E2) supplemented in the drinking water. Mice were allowed to drink and eat ad libitum for the duration of the study. Mice were weighed weekly as weight loss >10% was used as an indicator of acute toxicity. For tumor challenge, tumor cells were diluted in a 50/50 mixture of HBSS and phenol red-free LDEV-free Matrigel (Corning; Corning NY, USA). Mice were anesthetized with isoflurane, abdomen shaved, and challenged with 5×10^6^ MM134 or 44PE cells in the bilateral #4 mammary fat pads. Treatment was initiated when >25% of tumors exceeded ∼100mm^3^, with fulvestrant (5 mg/kg, diluted in 10% ethanol and 90% peanut oil, administered subcutaneously weekly), talazoparib (0.33 mg/kg, diluted in water with 4% DMSO + 30% PEG300 (Sigma-Aldrich), administered by oral gavage daily excluding weekends), or estrogen withdrawal (E2 supplement removed from drinking water; E2 supplementation was restored at the end of the indicated treatment window). Tumors were measured twice weekly using calipers, and mice were weighed weekly; tumor volume (*V*) was calculated using the formula *V* = (*S*^2^ x *L*) / 2, where *L* = Longest tumor diameter, and *S* = shortest tumor diameter. Mice were euthanized when any tumor diameter reached 15mm as defined humane endpoints.

Viable cryopreserved fragments of PDX HCI-013 were obtained from the Preclinical Research Resource at the University of Utah Huntsman Cancer Institute. PDX tumors were maintained in NSG mice with 30µM E2 supplemented in the drinking water. As above, naïve recipient NSG mice were maintained on E2 water for 1 week prior to PDX transplantation. For transplant, tumor was excised from donor mice, chopped into ∼1mm^3^ fragments, and held in HBSS to await transplant. Recipient mice were anesthetized with isoflurane, abdomen shaved, then a small incision was made near the #4 nipples for a single fragment of PDX tissue implanted into the exposed fat pat, with the wound closed with surgical staples. Buprenorphine-SR/ER (Wedgewood Pharmacy; Swedesboro, NJ 08085) was administered for analgesia. PDX were implanted bilaterally. Mice were treated as described above.

### Software

Statistical analyses and associated graphs were prepared with Graphpad Prism v10 (GraphPad Software, Boston, MA, USA).

---

**Supplemental Figure 1.**
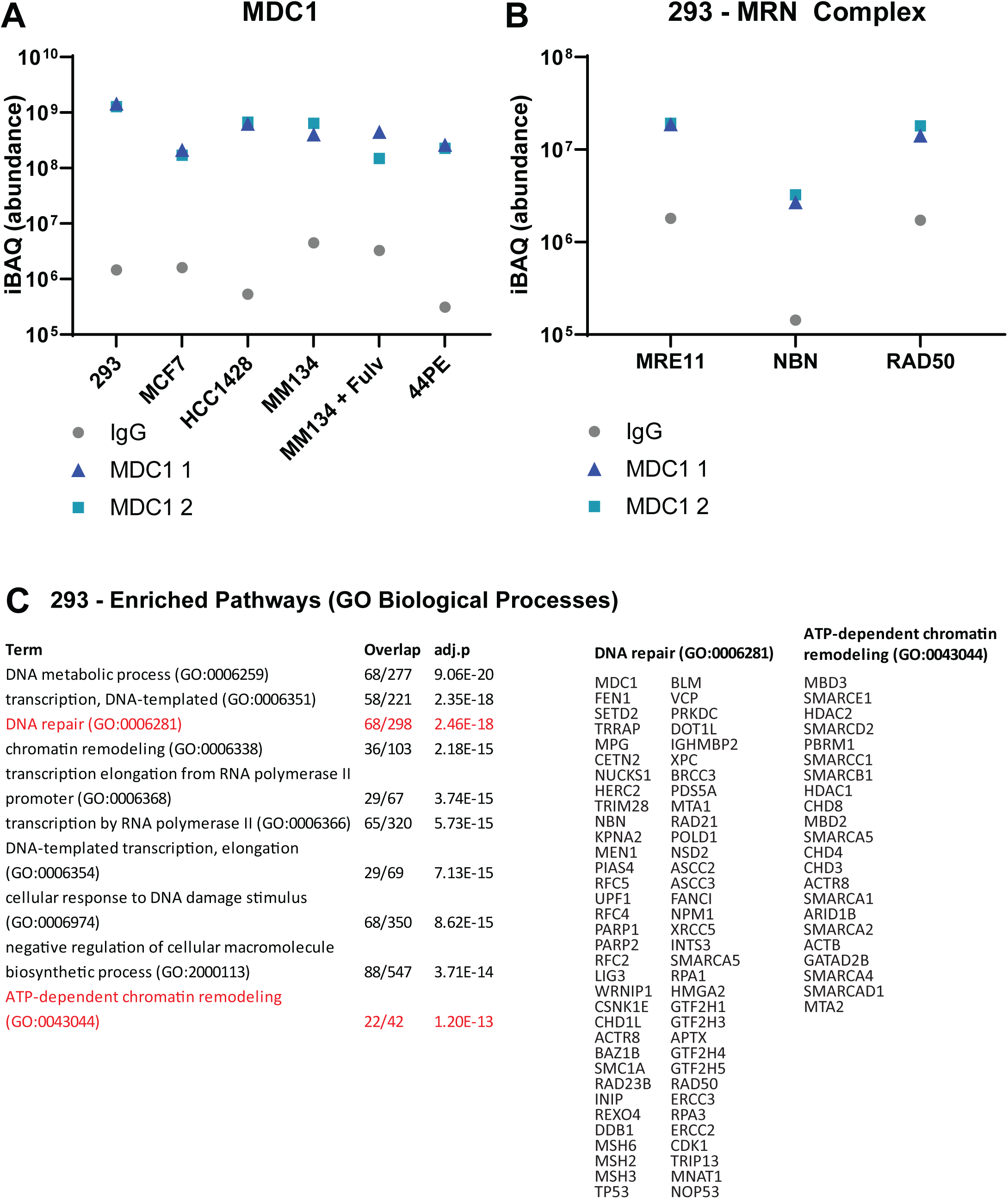
MDC1 is strongly enriched in IP/MS and key DNA repair partners are identified in 293 cells. **(A)** iBAQ abundance values for MDC1 peptides for biological duplicate MDC1 IP and IgG controls. **(B)** iBAQ abundance values from 293 cells, for MRN complex proteins as positive control for identifying previously reported constitutive MDC1 partners in 293 cells. **(C)** Gene set enrichment against GO Biological Processes, with identified proteins in two highlighted pathways at right.

**Supplemental Figure 2.**
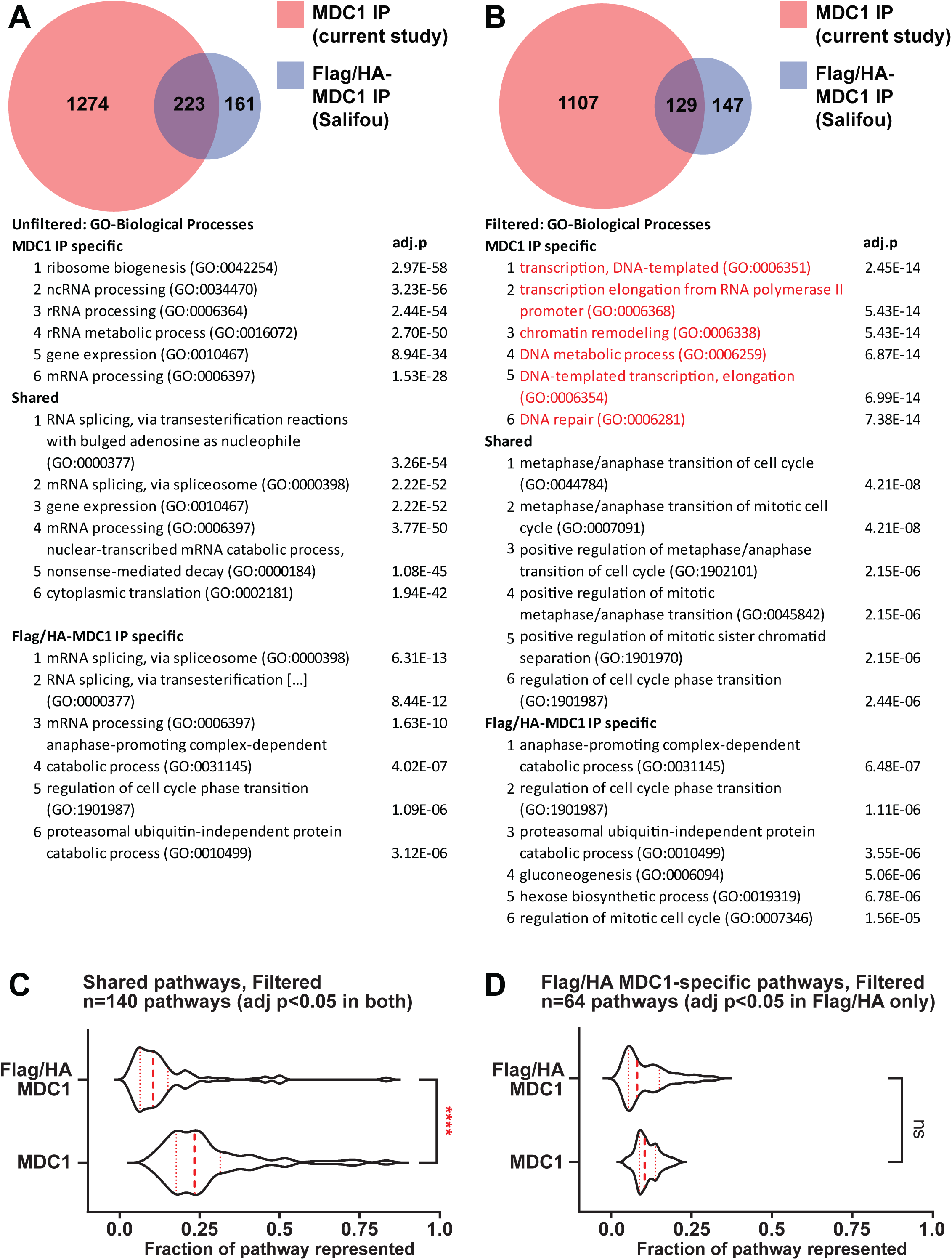
MDC1 IP/MS with specific antibody provided greater depth in pathway coverage and new target compared to affinity tag IP/MS. (A-B) Comparing 293 MDC1 IP/MS versus hits reported by Salifou et al (see text). **(A)** Gene set enrichment against GO Biological Processes, without filtering hits from RNA metabolic pathways, from study-specific versus shared MS hits. **(B)** Gene set enrichment against GO Biological Processes, after filtering hits from RNA metabolic pathways. DNA repair and associated pathways were among the top hits only from MDC1 antibody IP/MS. **(C)** MDC1 IP/MS provided greater pathway coverage among GO BP pathways enriched in both IP/MS studies. ****, t-test, p<0.0001. **(D)** For pathways that were enriched specifically in Salifou et al affinity tag IP/MS, MDC1 IP/MS provided equivalent pathway coverage in identified peptides.

**Supplemental Figure 3.**
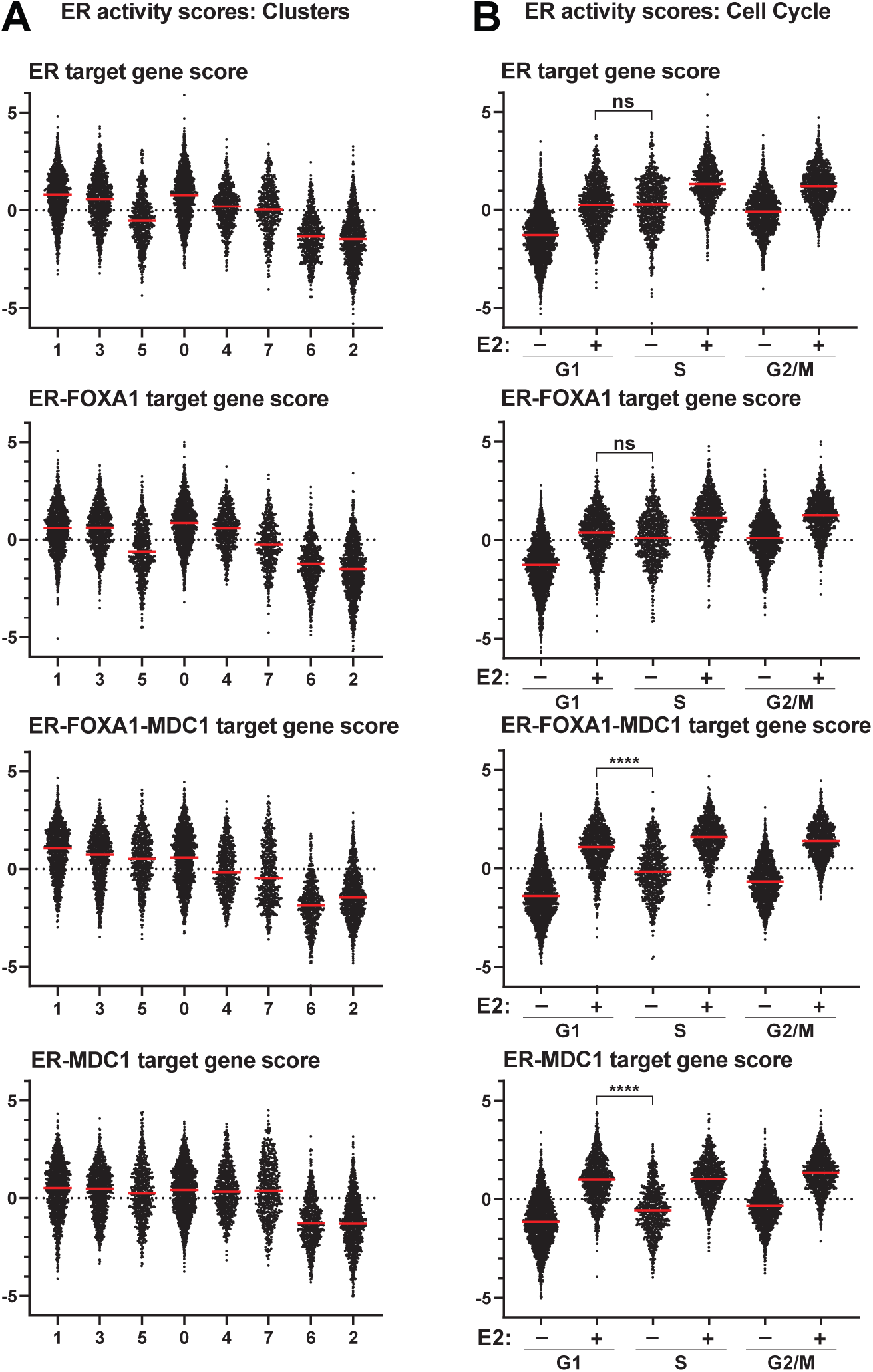
ER target gene sets and FOXA1 vs MDC1 activity are distinct across gene expression clusters, cell cycle status, and treatment group. **(A)** Single cell data represented by the heatmap in Figure 3C, red line = median score, points = individual cells. **(B)** ER target gene set scores by treatment (±E2) and predicted cell cycle state. Comparison between G1/+E2 and S/–E2 is representative of the cell cycle-dependent increase in non-MDC1 target genes (ER, ER-FOXA1 targets) compared to MDC1 target genes (ER- FOXA1-MDC1, ER-MDC1); i.e. for the former, target gene scores are increased in S/G2 relative to G1 regardless of estrogen treatment. ****, ANOVA, adj.p <0.0001.

**Supplemental Figure 4.**
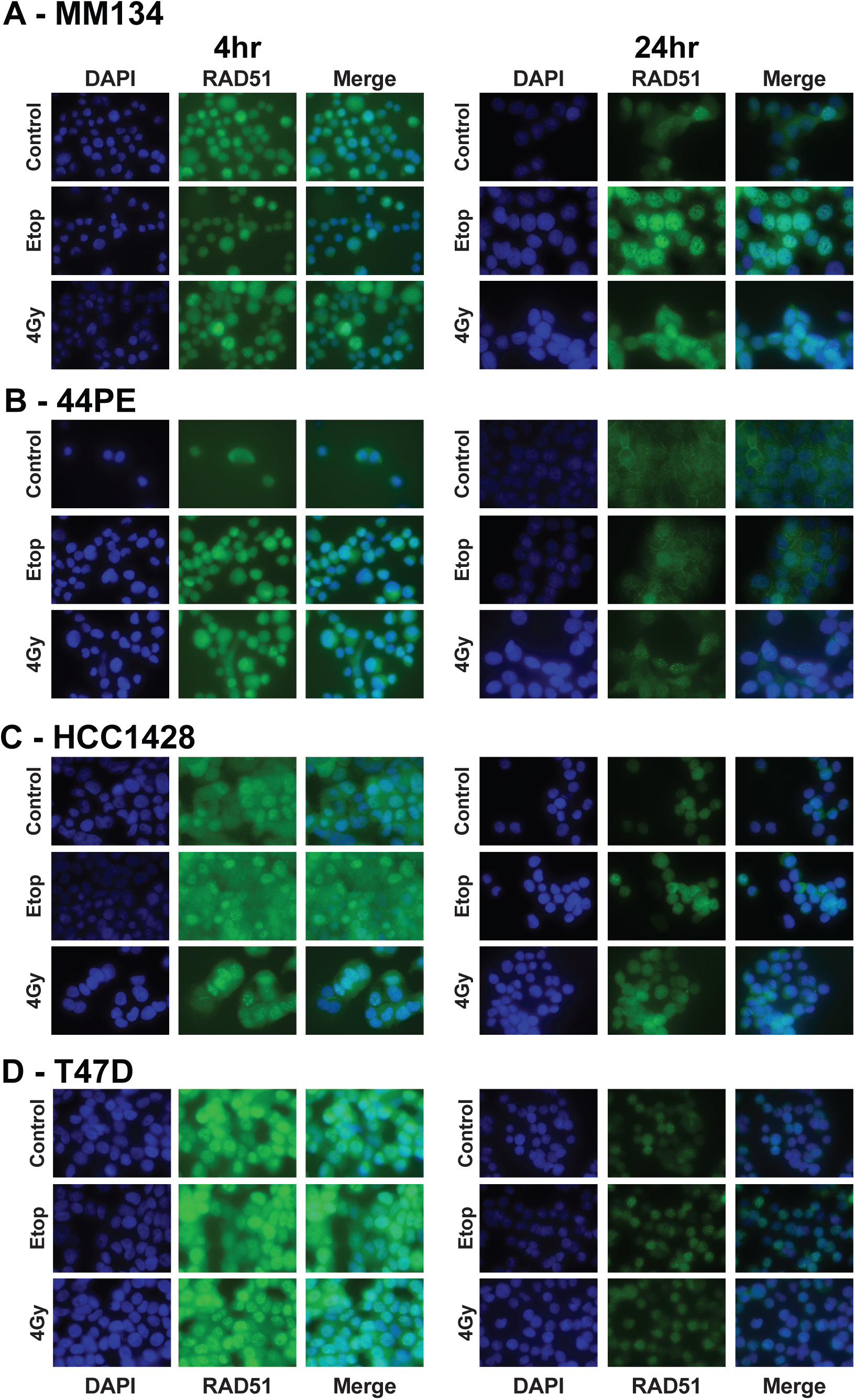
RAD51 foci formation in response to etoposide or XRT is delayed in ILC versus IDC cell lines. Representative immunofluorescent images of MM134 **(A)**, 44PE **(B)**, HCC1428 **(C)**, and T47D **(D)** cell lines in response to 10µM etoposide (Etop) or 4Gy of XRT at time points as described. Nuclear stain (DAPI, blue) and RAD51 (Green) are depicted individually and merged.

**Supplemental Figure 5.**
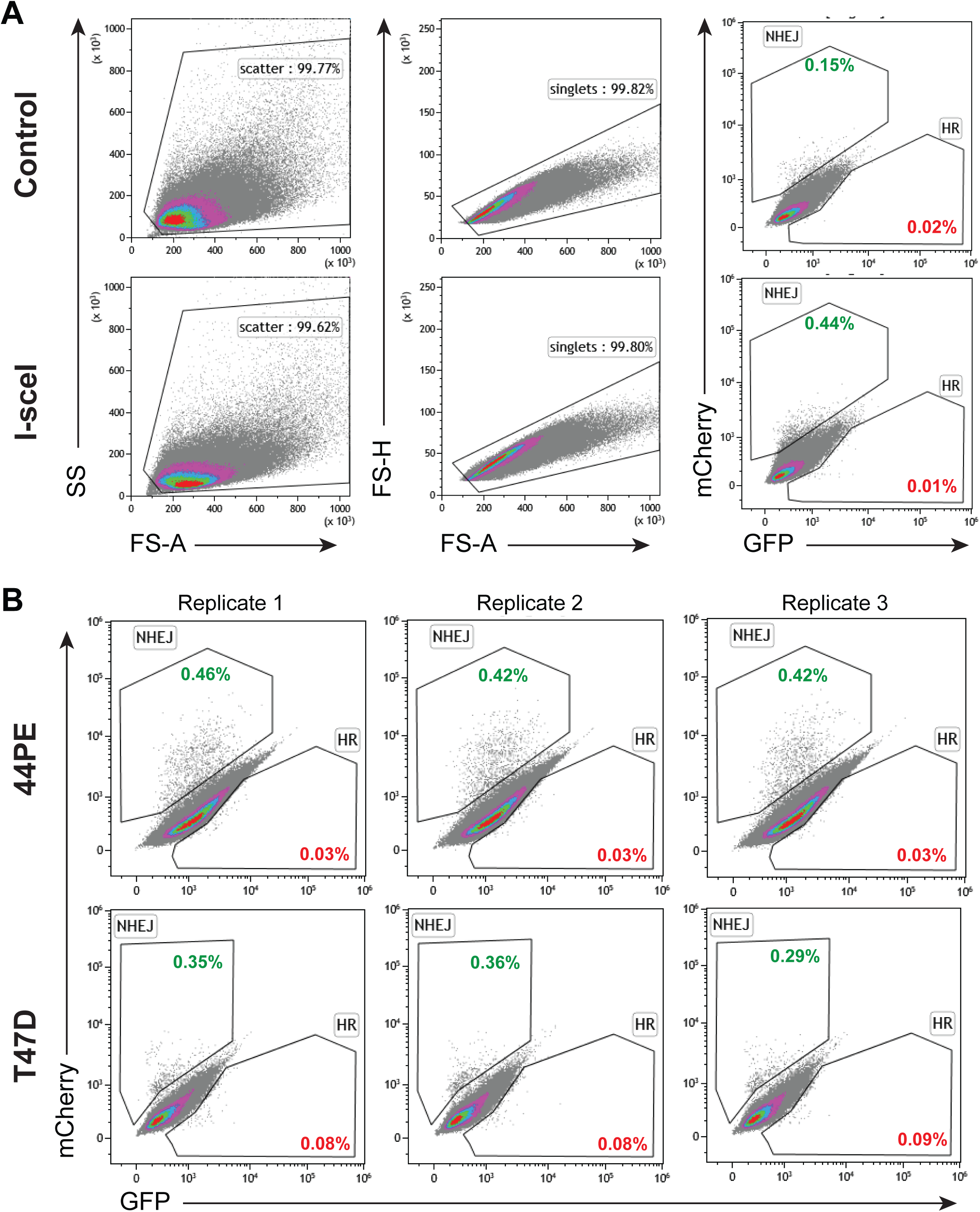
Gating strategy and representative data of stable TrafficLight Reporter integration. **(A)** Gating strategy of MM134 cells without (Control) and with (I-SceI) expression responses; percentages of gated events. **(B)** Representative FACS plots (n=3/line) of 44PE and T47D repair of the Traffic Light Reporter plasmid; percentages of gated events shown.

**Supplemental Figure 6.**
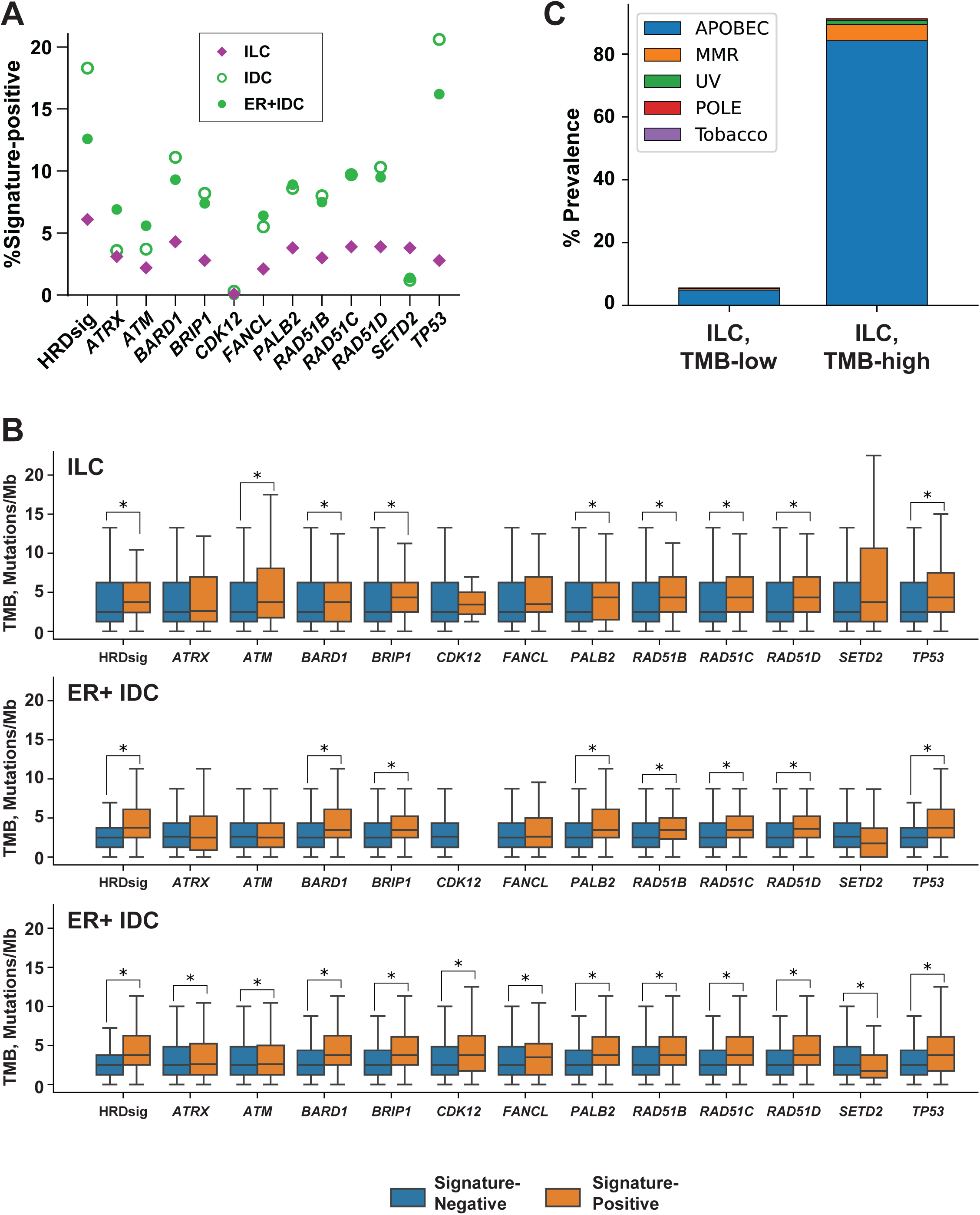
Mutational signature analyses in Foundation Medicine tumor cohort. **(A)** Relative frequency of the indicated scarring (HRDsig) and loss-of-function mutational signatures per gene. Relative differences between ILC and ER+ IDC for all signatures (except *CDK12*) are significant (chi-test p<1×10^-5^); *CDK12* signature is positive in only 55/22864 samples (0.24%). **(B)** TMB based on mutational signature status.*, Mann-Whitney U-test with multiple testing correction, adj.p<0.05. **(C)** Trinucleotide mutational signatures for ILC tumors. >80% of TMB-high ILC have an APOBEC signature, as described previously (24).

**Supplemental Figure 7.**
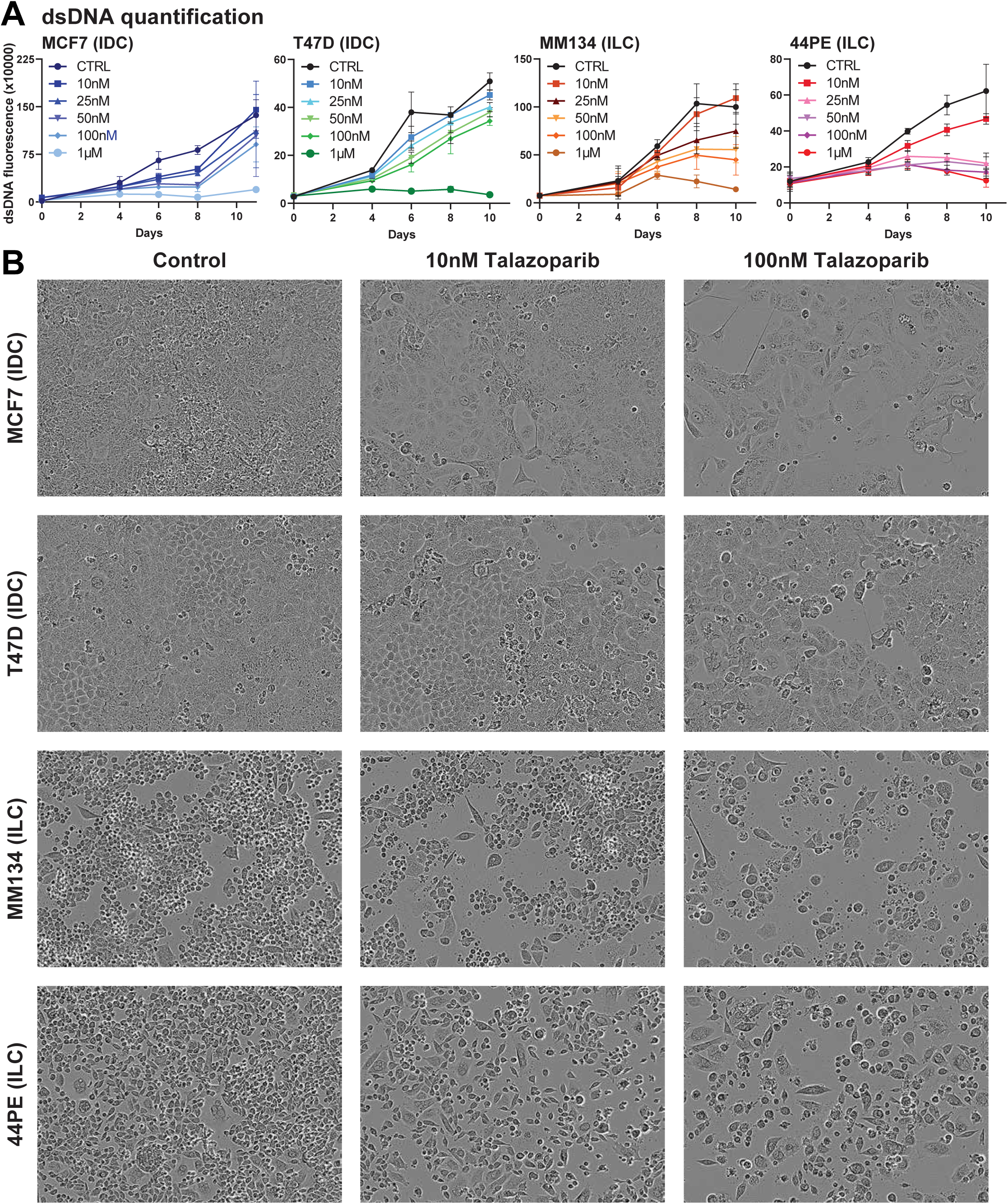
ILC cells are sensitive to PARPi talazoparib *in vitro*. **(A)** Time course of cell proliferation, assessed by dsDNA quantification, with increasing concentration of talazoparib. Extended >7d suggest decreased long-term viability in ILC cells specifically after the single talazoparib treatment. **(B)** Representative Incucyte images after 9 days of treatment as indicated. Remaining IDC cells are fewer than control but morphologically appear healthy and viable, whereas ILC cells show irregular morphology consistent with apoptosis or other cell death.

**Supplemental Figure 8.**
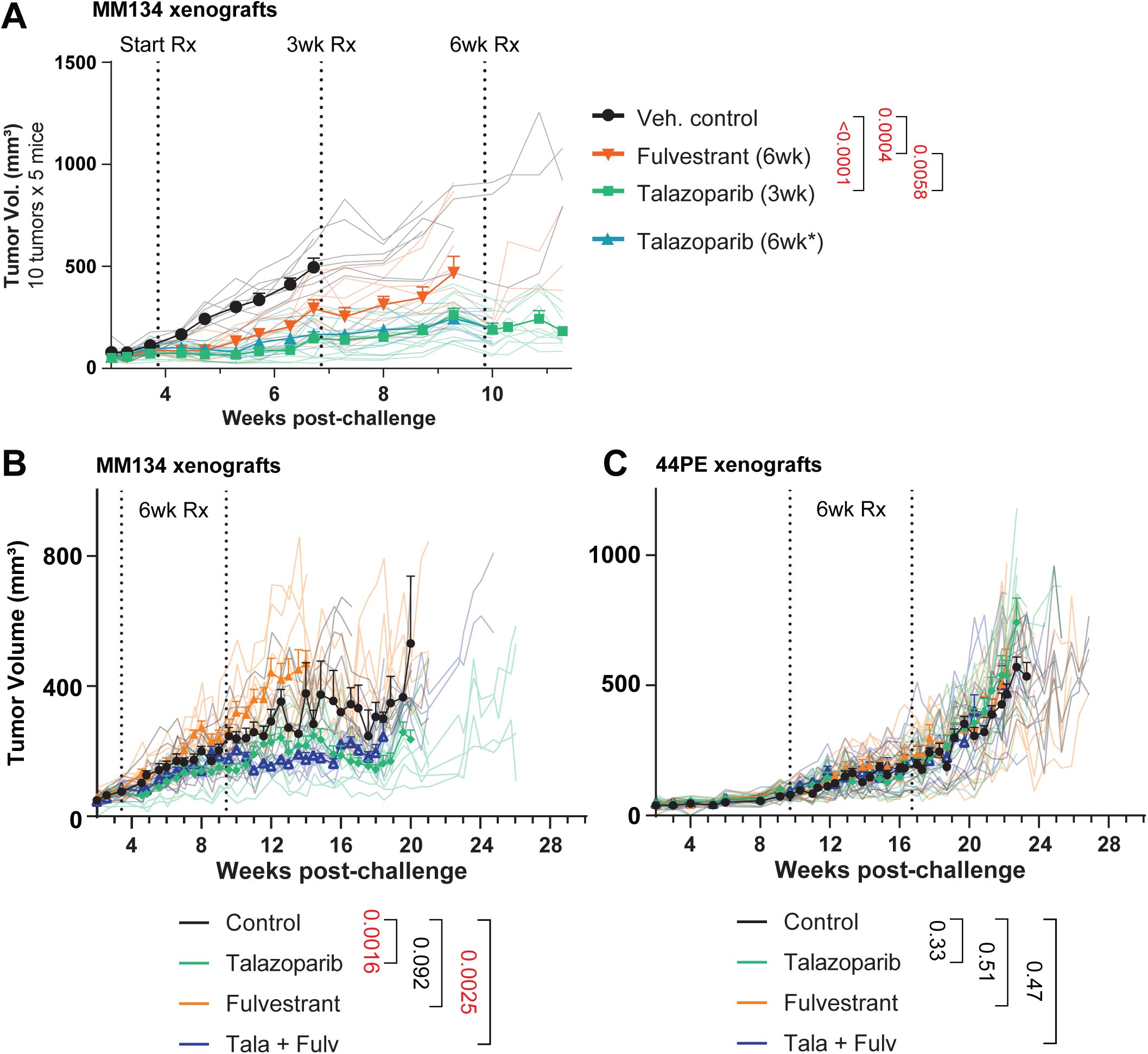
ILC cells are sensitive to PARPi talazoparib *in vivo*. Bold lines/symbols show mean tumor size ± SEM; individual tumor size shown as matching faded lines. Bold lines ended at first tumor size human endpoint reached per arm. Two-way ANOVA treatment effect p-values shown. **(A)** 6wk talazoparib data excluded from statistical analyses. **(B-C)** Control and Talazoparib arms also shown in Figure 7.

**Supplemental Figure 9.**
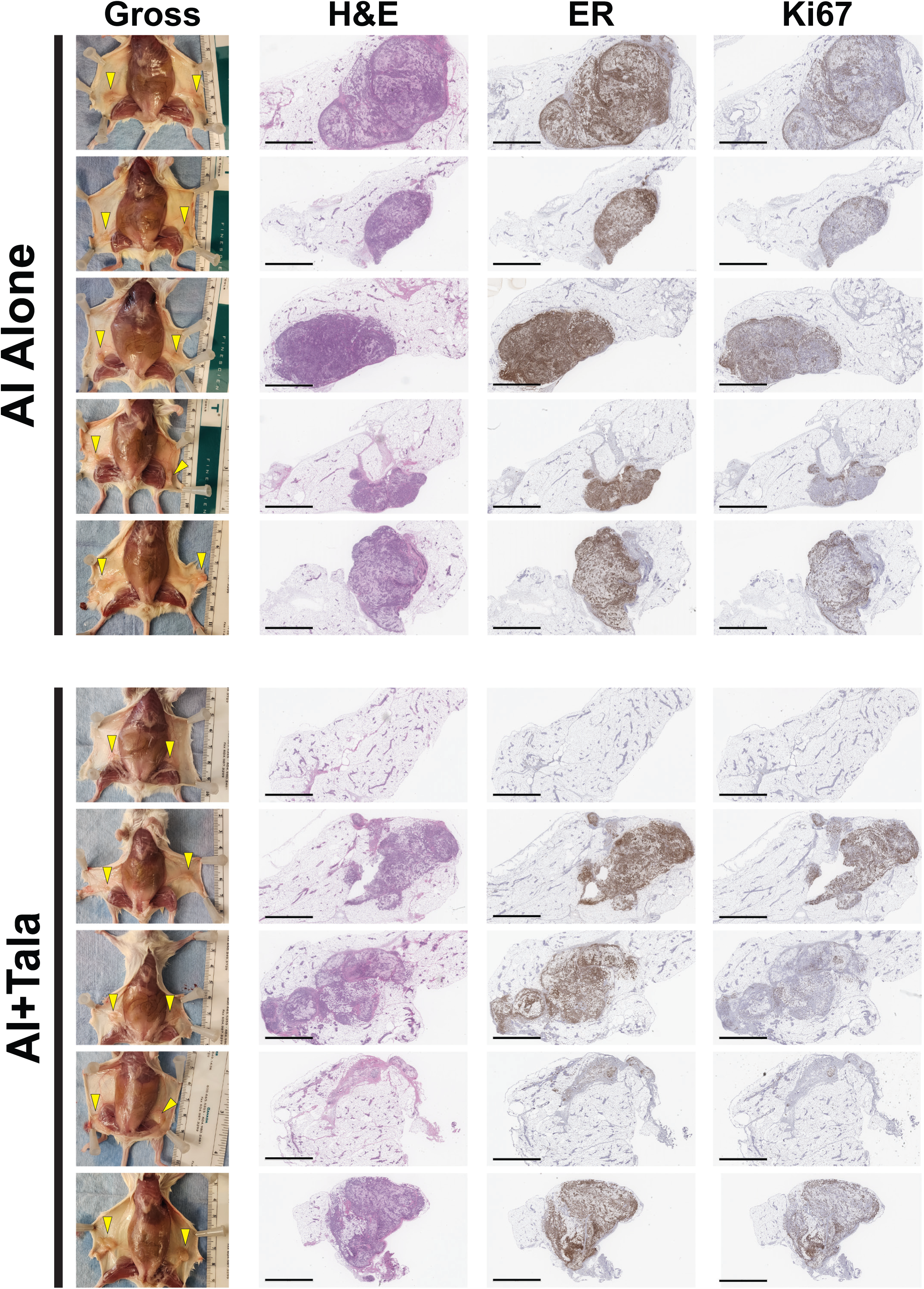
Necropsy and histology images for HCI-013 at 10 week treatment completion. Mice were sacrificed and treatment completion to examine on-treatment tumor phenotype; all control and tala- alone mice were removed from study earlier than 10wks based on humane tumor size endpoints. At left, gross images at necropsy show tumor beds (yellow arrows). Several mice in the AI+Tala cohort with non-palpable tumors had residual masses visible at necropsy, but had limited ILC cellularity by histology (at right).

**Supplemental Figure 10.**
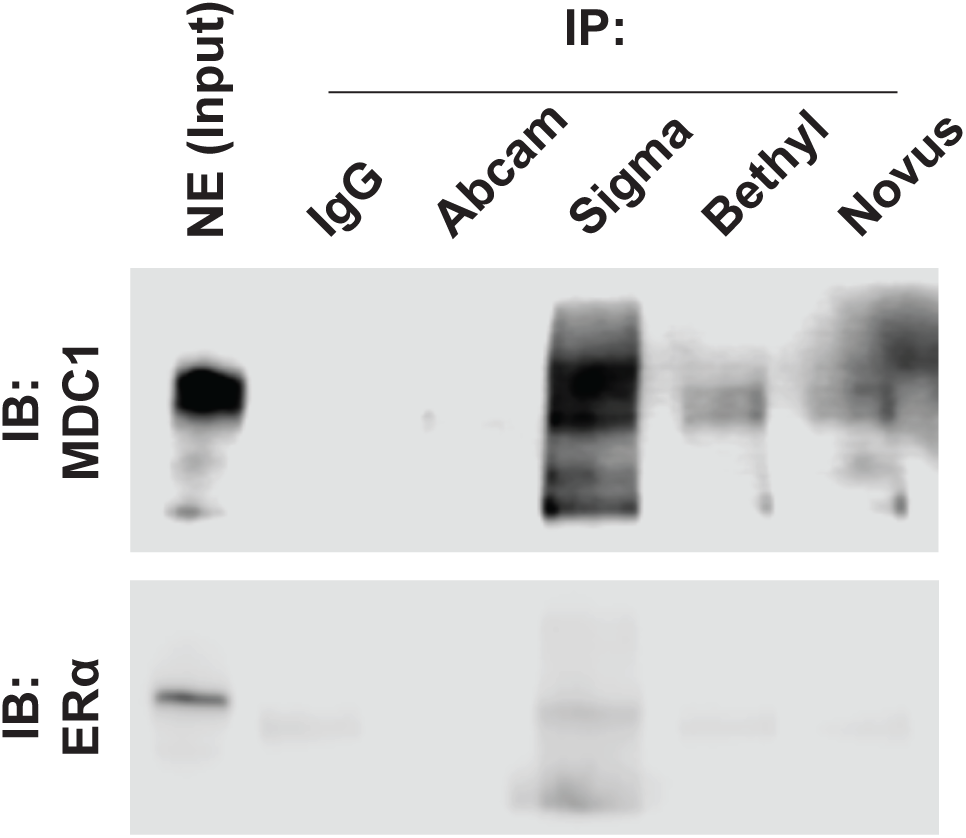
Identification and optimization of MDC1 antibodies for use in co-immunoprecipitation. Comparison of Immunoprecipitation (IP) with 5µg of: IgG (Jackson ImmunoResearch), Abcam (ab11171), Sigma (M2444), Bethyl (A300-051A), and Novus (NB 100-395) anti-MDC1 antibodies. Immunoblot was probed with Sigma (M2444) or ESR1/ERα (6F11). Sigma-M244 had the greatest enrichment compared to IgG and the greatest co-IP of ESR1.

